# Sphingosine 1-phosphate receptor-1 signaling enhances neutrophil survival while suppressing inflammation and bacterial host defense functions

**DOI:** 10.64898/2026.02.05.703783

**Authors:** Yueh-Chien Lin, Takahiro Seno, Alan Y. Hsu, Andréane Cartier, Andrew Kuo, Michel V Levesque, Avishek Ghosh, Ingo Fohmann, Victoria A. Blaho, Sylvain Galvani, Reed Crocker, Samuel W Kazer, José Ordovás–Montañés, Hongbo R. Luo, Timothy Hla

## Abstract

Sphingosine 1-phosphate (S1P), a lipid mediator that signals through five G protein-coupled S1P receptors (S1PRs), regulates T cell trafficking, tissue residency, and inflammatory processes. In contrast to the established roles of S1P signaling in T cell trafficking, its role in neutrophil biology remains poorly understood. Here, we demonstrate that S1PR1, one of the two S1PRs expressed in neutrophils, promotes mitochondrial fitness, enhances survival, and reduces inflammatory output. Using myeloid-and neutrophil-selective S1PR1 overexpression (S1PR1^hi^) mouse models, we show that elevated S1PR1 signaling promotes redistribution of neutrophils from the bone marrow to peripheral tissues under steady-state conditions, without inducing overt inflammation or tissue injury. S1PR1^hi^ neutrophils display altered surface marker profiles consistent with a less mature state. These cells also exhibit reduced *in vivo* turnover, increased mitochondrial membrane potential and oxidative phosphorylation, and transcriptional programs linked to survival and dampened inflammatory signaling. Functionally, S1PR1^hi^ neutrophils exhibit a reduced oxidative burst while preserving phagocytic capacity. However, *in vivo* bacterial challenge revealed impaired bacterial clearance in the lung. In contrast, in a model of influenza A virus infection of the lung, enhanced neutrophil-intrinsic S1PR1 signaling conferred reduced lung injury, decreased inflammatory output, and improved survival. Together, these findings support a model in which S1PR1 reprograms neutrophils, enabling their survival and dampening inflammatory potential in a context-dependent manner, thereby differentially shaping host defense and tissue protection during microbial infections.

**One Sentence Summary:** S1PR1 signaling reprograms neutrophils into a persistent, low-inflammatory state that confers protection against viral lung injury but impairs antibacterial host defense.

## INTRODUCTION

Neutrophils are the primary circulating innate immune cells and play crucial roles in the inflammatory response (1, 2). Following infection or tissue injury that increases vascular permeability, neutrophils rapidly extravasate into affected tissues, where they initiate inflammation by secreting lipid mediators, reactive oxygen species, chemokines, cytokines, and proteases. These events drive and amplify the early stages of inflammation, facilitating the removal of infectious or noxious agents and allowing subsequent mechanisms of tissue repair and the restoration of homeostasis to ensue (3). Neutrophils mediate phagocytosis, release reactive oxygen species (ROS), defensins, and lytic enzymes, and form neutrophil extracellular traps (NETs), all of which are critical for inflammation (4). While these mechanisms are essential for host defense against pathogens, they also contribute to sterile inflammation and, in some contexts, to carcinogenesis and tumor progression (5), unregulated neutrophil-derived mechanisms can also lead to collateral tissue damage during sterile inflammation (6, 7) and to atherothrombotic and autoimmune diseases (8).

Beyond their well-established roles in initiating inflammation, neutrophils are increasingly recognized for their contributions to the resolution phase of inflammation, a mechanistically programmed event necessary for the return to physiological homeostasis (9, 10). In this context, neutrophil elaboration of specialized pro-resolving lipid mediators (SPMs) and their responses have been characterized in numerous models of inflammation resolution and host defense (11–14). Known SPMs are largely derived from the oxidative metabolism of omega-3 polyunsaturated fatty acids, namely docosahexaenoic acid, eicosapentaenoic acid, and linolenic acid. These SPMs are thought to interact with GPCRs on cells in a stereospecific manner to regulate processes critical for the resolution of inflammation, such as phagocytosis and the production of anti-inflammatory factors. Whether other lipid mediators derived from the metabolism of membrane phospholipids, particularly those signaling through GPCRs, regulate neutrophil resolution programs remains unclear.

Recent studies have shown that neutrophils display marked heterogeneity, with distinct phenotypes defined by transcriptional, functional, and developmental programs (15–20). Recent work has further refined neutrophil classification frameworks, highlighting functional diversity across developmental and tissue-associated states (21). These subsets participate in a multitude of physiological processes, including inflammation resolution (22, 23), tissue remodeling (24, 25), angiogenesis (26, 27), and regulation of hematopoiesis (3, 28). Despite this functional diversity, the upstream signals that control neutrophil phenotypic plasticity and constrain inflammatory potential remain poorly understood.

Although neutrophil development is classically viewed as a bone marrow–restricted process, accumulating evidence indicates that the spleen serves as an important peripheral reservoir and regulatory niche for neutrophils (29, 30). Splenic neutrophils differ from bone marrow counterparts in maturation state, trafficking behavior, and functional programming, and can contribute to both host defense and the resolution of inflammation. Extramedullary myelopoiesis in the spleen can also occur under homeostatic and inflammatory conditions, providing a site for ongoing neutrophil differentiation and functional tuning. Moreover, neutrophils appear to undergo continued maturation and phenotypic adaptation after egress from the bone marrow, influenced by tissue-specific cues within secondary lymphoid organs such as the spleen (29, 31–33). These splenic microenvironments expose neutrophils to distinct cytokine, chemokine, and lipid mediator landscapes that shape their survival, trafficking, and inflammatory potential. However, specific lipid-sensing pathways that mediate these tissue-specific adaptations remain largely undefined, and differential expression of trafficking receptors and signaling molecules may reflect functional specialization rather than a simple developmental stage.

Sphingosine-1-phosphate (S1P) is a bioactive lipid mediator generated during membrane sphingolipid metabolism. It signals through five G protein-coupled receptors (GPCRs), now known as S1PR1-5 (34, 35), and plays crucial roles in lymphocyte trafficking, vascular development, and immune homeostasis (36, 37). S1P is best known for regulating lymphocyte egress from the thymus and secondary lymphoid organs into the circulation (38). S1PR1 functional antagonists that preferentially activate β-arrestin have been approved for the treatment of autoimmune diseases, underscoring the importance of adaptive immune cell S1PR1 signaling in autoimmune pathology (37, 39). More recently, S1P has been shown to influence T cell metabolism (40) and differentiation fate, including the balance between Th17 and regulatory T cells (41–43). However, the role of S1P receptors in myeloid lineage cells, particularly neutrophils, remains incompletely understood. Prior studies suggest that S1P can influence neutrophil functions, such as pain sensitization (hyperalgesia) (44) and fungus-induced vasculitis (45), while genetic disruption of S1P synthesis by knocking out sphingosine kinases does not markedly impair neutrophil migration in acute inflammatory models (46). Elevated systemic S1P levels in S1P lyase-deficient mice result in neutrophilia that is reversed by deletion of S1PR4, implicating receptor-specific S1P signaling in neutrophil biology (47). A rare missense mutation in *S1PR4* has also been linked to human neutropenia (48). Although these findings support a role for S1P receptors in neutrophil regulation, they do not directly address the contribution of S1PR1 to neutrophil homeostasis or functional state. Notably, apolipoprotein M (ApoM)-bound S1P suppresses NETosis via activation of S1PR1 and S1PR4, suggesting receptor-specific anti-inflammatory functions (49). However, whether S1PR1 signaling is critical for neutrophil homeostasis, functions (inflammation, resolution), and heterogeneity remains unclear. Specifically, the direct contribution of S1PR1, a therapeutic target in many autoimmune diseases with five FDA-approved drugs in the clinic, to neutrophil trafficking, survival, phenotype, or function under inflammatory or infectious conditions remains unknown. Given the clinical use of S1PR modulators, understanding their effects on innate immunity is warranted.

In this study, we used myeloid-and neutrophil-restricted S1PR1 gain-and loss-of-function mouse models to define the role of S1PR1 signaling in neutrophil biology. We show that sustained S1PR1 signaling reprograms neutrophils into a long-lived, metabolically supported state characterized by reduced oxidative and inflammatory responses and altered surface marker profiles consistent with a distinct neutrophil state. Functionally, S1PR1^hi^ neutrophils retain intrinsic antibacterial activity under simplified *ex vivo* conditions, but exhibit impaired bacterial clearance *in vivo*, particularly in the lung, while also conferring enhanced protection against viral infection-induced lung injury. Together, our findings identify S1PR1 as a key regulator of neutrophil state that tunes the balance between host defense and inflammatory tissue damage in a context-dependent manner. These results may have therapeutic implications for infectious, autoimmune, and inflammatory diseases.

## RESULTS

### Myeloid S1PR1 expands systemic neutrophil compartments under steady-state conditions

We generated two mouse models of S1PR1 loss-and gain-of-function: LysM*-Cre; S1pr1^f/f^*(LysM*-S1pr1* KO) and LysM*-*Cre; *S1pr1^flox-stop-flox^*(LysM*-S1pr1* TG). Both mouse strains were viable and appeared phenotypically normal. Complete blood count analysis revealed that LysM*-S1pr1* TG mice had elevated numbers of circulating white blood cells, neutrophils, and monocytes, with no significant changes in other hematopoietic lineages (Fig. 1, A–D). In contrast, LysM*-S1pr1* KO mice showed no significant changes in circulating neutrophil or monocyte numbers but displayed reduced lymphocyte counts and a modest decrease in total leukocyte counts (Fig. 1A–D). These data suggest that S1PR1 is not required for neutrophil and monocyte numbers in the circulation, but increased S1PR1 expression in the myeloid lineage is associated with expansion of the neutrophil and monocyte compartments in the blood. Reduced lymphocytes in LysM*-S1pr1* KO mice could reflect recombination occurring in early hematopoietic progenitors that transiently express Lyz2, rather than being restricted exclusively to mature myeloid cells (50, 51). LysM*-S1pr1* TG mice exhibited reduced red blood cell (RBC) count, hemoglobin levels, and hematocrit, along with increased mean corpuscular volume (MCV) and mean corpuscular hemoglobin (MCH) (Fig. S1, A–E), indicating that the TG mice had mild macrocytic normochromic anemia. Wild-type neutrophils from bone marrow and the spleen expressed *S1pr1* mRNA, and as expected, LysM*-S1pr1* TG neutrophils showed increased *S1pr1* mRNA expression (Fig. S1F). *S1pr1* mRNA levels were higher in splenic *vs.* BM neutrophils in both LysM*-S1pr1* TG and control mice, suggesting that S1PR1 expression is increased in peripheral neutrophils, which are more mature. We observed a modest increase in *S1pr4* transcript levels in neutrophils from *S1pr1* TG mice; however, the magnitude of this change was minor and more variable than the upregulation of *S1pr1* (Fig. S1G).

**Fig. 1.**
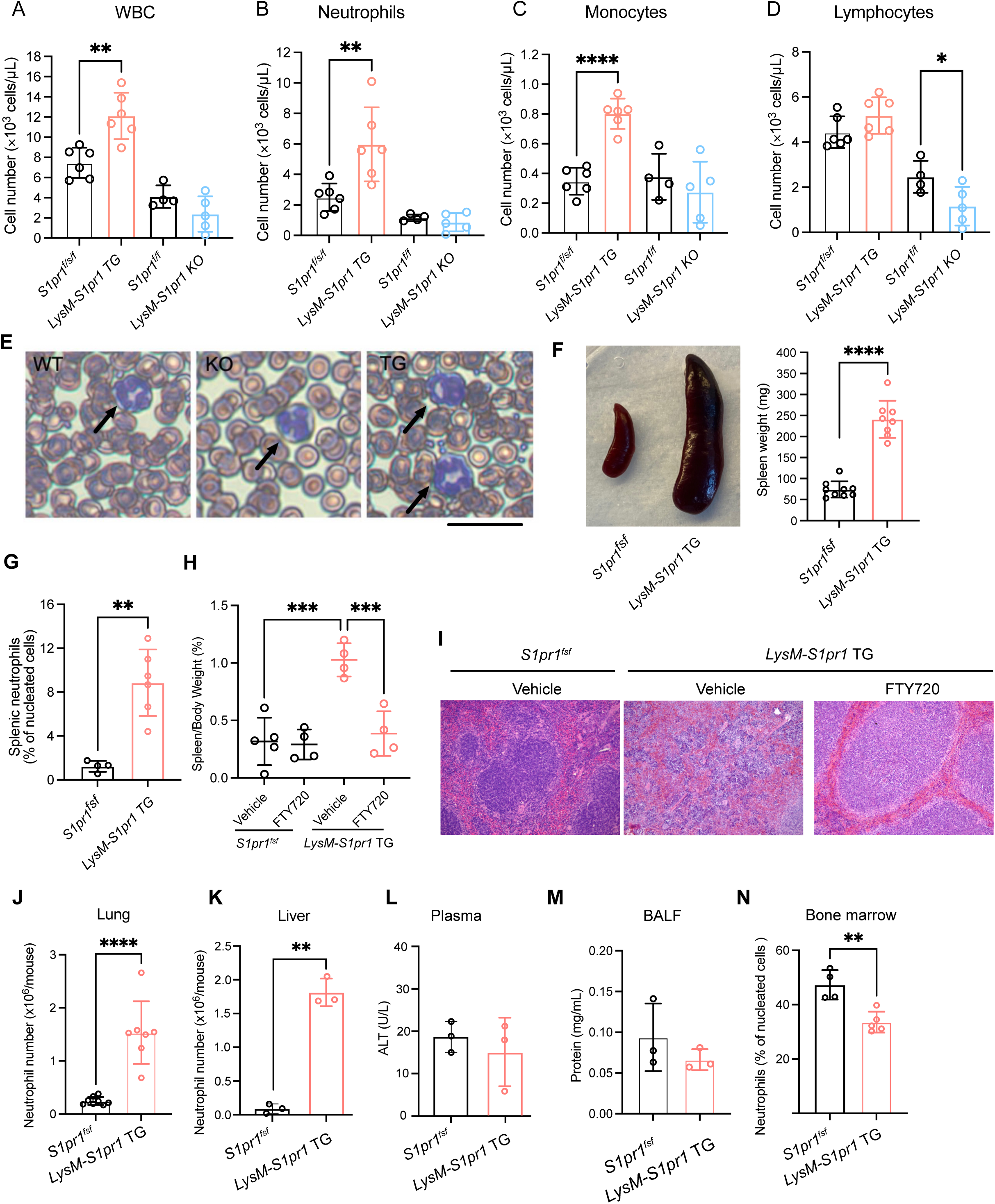
Myeloid S1PR1 is associated with systemic neutrophil expansion and redistribution. **(A-D)** Counts of white blood cells (WBCs), neutrophils, monocytes, and lymphocytes in peripheral blood from LysM*-S1pr1* TG (n = 6), *S1pr1^fsf^* (n = 6), LysM*-S1pr1* KO (n = 4), and *S1pr1^f/f^* (n = 5) mice. Total blood cell counts were measured using the HemaVet system. **(E)** Representative blood smear images of neutrophils in wild-type (WT), LysM*-S1pr1* KO, and TG mice. **(F)** Images and weights of spleens from *S1pr1^fsf^*(n = 9) and LysM*-S1pr1* TG (n = 8) mice. **(G)** Flow cytometric quantification of splenic neutrophil frequency (CD45^+^CD11b^+^Ly6G^+^) in *S1pr1^fsf^* (n = 4) and LysM*-S1pr1* TG (n = 6) mice. **(H)** Spleen/body weight ratio in vehicle-or FTY720-treated *S1pr1^fsf^* and LysM-*S1pr1* TG mice (n ≥ 4 mice per group). **(I)** Representative H&E images of spleens after FTY720 or vehicle treatment. **(J and K)** Neutrophil counts in the lung **(J)** (n ≥ 7 mice per group) and liver **(K)** (n = 3 mice per group). **(L)** Plasma alanine transaminase (ALT, U/L) level (n = 3 mice per group). **(M)** Total protein in BALF (n = 3 mice per group). **(N)** Bone marrow neutrophil frequency among WBCs in *S1pr1^fsf^*(n = 4) and LysM*-S1pr1* TG (n = 5) mice. Data are mean ± SD. \**p* < 0.05, \*\**p* < 0.01, \*\*\**p* < 0.001, \*\*\*\**p* < 0.0001 (one-way ANOVA with multiple comparisons, or two-tailed unpaired Student’s *t* test).

Given the elevated blood neutrophil counts in LysM*-S1pr1* TG mice, we next examined neutrophil morphology and distribution in hematopoietic organs. Circulating neutrophil morphology was unremarkable in both TG and KO mice (Fig. 1E). However, LysM*-S1pr1* TG mice exhibited marked splenomegaly, with spleens ∼3× larger than those of littermate controls (Fig. 1F). Flow cytometry revealed a >9-fold increase in splenic neutrophils (CD45^+^CD11b^+^Ly6G^+^) (Fig. 1G). In contrast, LysM*-S1pr1* KO mice showed no changes in spleen size or splenic neutrophil numbers (data not shown). To assess whether the splenomegaly phenotype is sensitive to pharmacologic S1PR inhibition, we treated LysM*-S1pr1* TG mice with FTY720, a functional antagonist of S1PR1, for 2 weeks. Spleen size returned to normal littermate control levels after FTY720 treatment (Fig. 1H), and H&E staining showed restoration of normal splenic architecture (Fig. 1I), indicating that the splenomegaly is induced by neutrophil S1PR1^hi^ states. In addition, resident neutrophils in the lung and liver were elevated in LysM*-S1pr1* TG mice (Fig. 1, J and K) without detectable signs of inflammation or damage to the lung or liver, such as high protein content in bronchoalveolar lavage fluid (BALF) or elevated alanine aminotransferase (ALT) enzyme activity in plasma (Fig. 1, L and M), respectively. Conversely, bone marrow neutrophils were significantly reduced in LysM*-S1pr1* TG mice (Fig. 1N), suggesting increased homeostatic mobilization of neutrophils into the S1P-rich circulatory compartment.

To determine whether the consequence of S1PR1 overexpression is neutrophil-intrinsic, we used a neutrophil-selective Mrp8-Cre driver to overexpress S1PR1 (Mrp8*-S1pr1* TG mice). Similar to LysM-Cre counterparts, Mrp8*-S1pr1* TG mice displayed increased circulating neutrophil counts, enlarged spleens, elevated spleen/body weight ratio and splenic neutrophil frequency, and reduced bone marrow neutrophil percentage and count (Fig. S2), supporting a cell-intrinsic role for S1PR1 signaling. These findings suggest that S1PR1^hi^ neutrophils show increased egress from the bone marrow, circulate longer, and reside in peripheral organs without causing organ damage under homeostatic conditions. In contrast, lack of S1PR1 in neutrophils did not affect steady-state granulopoiesis, circulating neutrophil numbers, or tissue residence, suggesting that this pathway is not required for general neutrophil egress from the bone marrow, survival, and functions *in vivo*.

### Sustained S1PR1 signaling alters neutrophil phenotypes

Flow cytometry showed that surface S1PR1 levels were low in both TG and control splenic neutrophils, whereas total S1PR1 signal was markedly higher in LysM*-S1pr1* TG neutrophils (Fig. 2A). ImageStream cytometry analysis showed that S1PR1 was predominantly intracellular, a pattern consistent with ligand-induced S1PR1 endocytosis (Fig. 2B). Given that sustained GPCR activation is commonly associated with receptor endocytosis, these data suggest ligand-induced S1PR1 internalization and sustained receptor engagement in TG neutrophils.

**Fig. 2.**
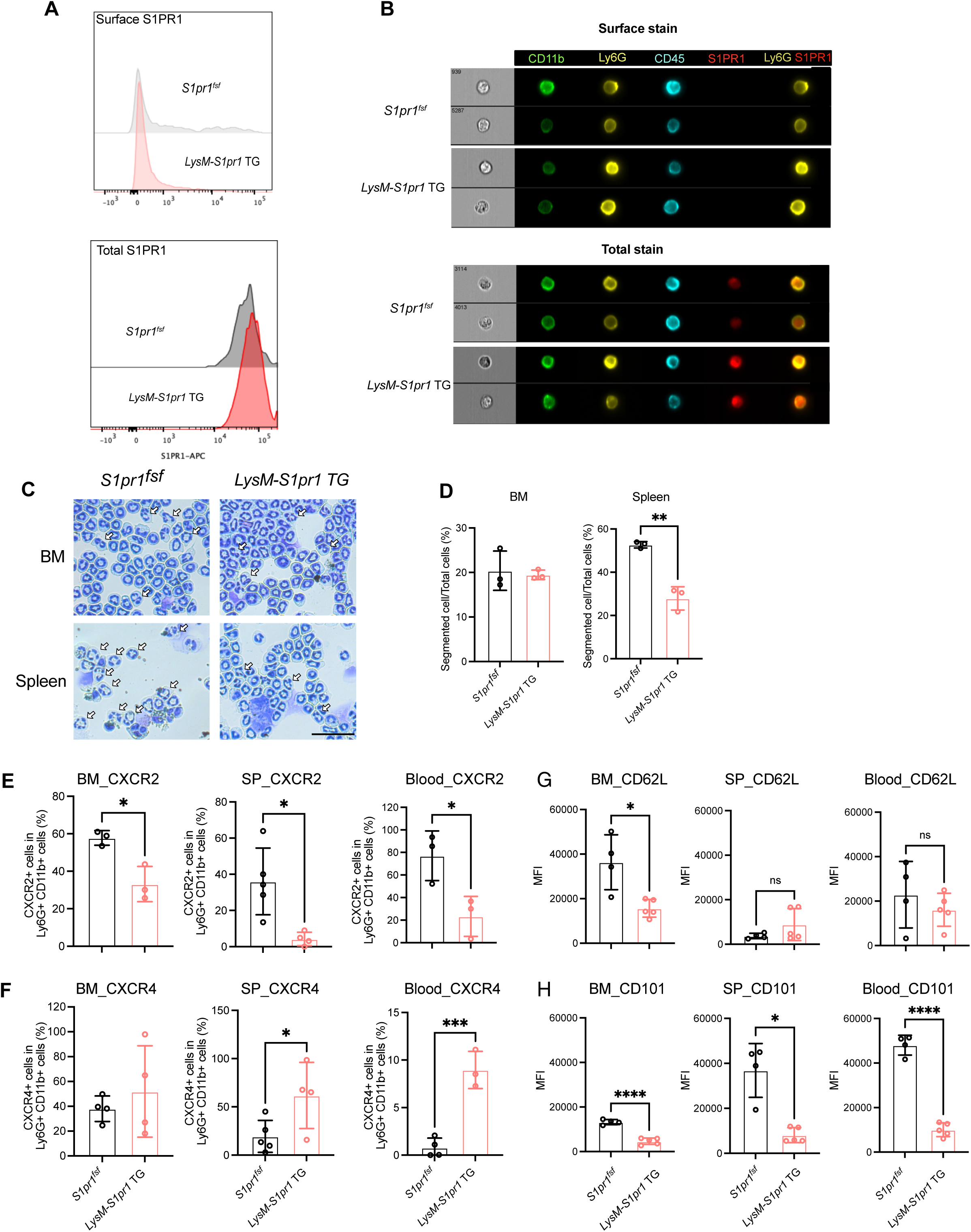
Sustained S1PR1 signaling alters neutrophil phenotypes. **(A)** Flow cytometric analysis of surface and total S1PR1 staining on splenic neutrophils (n ≥ 3 mice per group). **(B)** ImageStream analysis showing intracellular localization of S1PR1 in individual splenic neutrophils (n ≥ 3 mice per group). **(C)** Giemsa staining of isolated bone marrow (BM) and splenic (Spleen) neutrophils; segmented nuclei indicated by white arrows. Scale bar = 20 μm. **(D)** Quantification of segmented nuclei in BM and splenic neutrophils (n = 3 mice per group). **(E and F)** Flow cytometric analysis of CXCR2^+^ **(E)** and CXCR4^+^ **(F)** neutrophils in BM (n ≥ 4 mice per group), spleen (n ≥ 4 mice per group), and blood (n ≥ 3 mice per group). **(G and H)** Mean fluorescence intensity (MFI) of CD62L **(G)** and CD101 **(H)** on neutrophils (CD11b+Ly6G+) in BM, spleen, and blood. Quantification on the right. Data are mean ± SD. \**p* < 0.05, \*\**p* < 0.01, \*\*\**p* < 0.001, \*\*\*\**p* < 0.0001 (two-tailed unpaired Welch’s *t* test).

Giemsa staining of splenic neutrophils revealed an increased frequency of neutrophils with band-shaped nuclei and reduced segmentation in LysM*-S1pr1* TG mice (Fig. 2, C and D), consistent with phenotypes commonly associated with less mature neutrophil states (52–54). However, no apparent change was observed in the bone marrow, which contains newly formed neutrophils. Additionally, surface CXCR2 was downregulated and CXCR4 upregulated in LysM*-S1pr1* TG neutrophils across bone marrow, spleen, and blood (Fig. 2, E and F), a pattern previously associated with altered neutrophil trafficking and post–bone marrow adaptation. To further characterize the phenotypic state of S1PR1^hi^ neutrophils, we assessed surface markers associated with neutrophil maturation and activation. CD62L expression was largely preserved across compartments, with only a modest reduction in bone marrow neutrophils (Fig. 2G), suggesting that S1PR1^hi^ neutrophils do not exhibit classical activation-associated CD62L shedding. In contrast, CD101 expression was markedly reduced, particularly in bone marrow and circulating neutrophils (Fig. 2H), indicating a shift toward a distinct neutrophil state. Together, these data indicate that S1PR1^hi^ neutrophils display altered phenotypic features consistent with modified maturation and trafficking states.

To determine whether S1PR1 overexpression influences myelopoiesis and/or granulopoiesis, we profiled myeloid subsets in the spleen and bone marrow by flow cytometry. Megakaryocyte/erythrocyte progenitors (MEPs) were significantly increased in the spleen (Fig. S3), consistent with compensatory erythroid responses. Common myeloid progenitors (CMPs) were mildly elevated, whereas granulocyte myeloid progenitors (GMPs) remained unchanged. Bone marrow progenitor frequencies were slightly elevated across all subsets. Together, these data indicate that increased S1PR1 signaling in neutrophils regulates its phenotype, characterized by CXCR4^hi^, CXCR2^lo^, with increased circulatory transit time and residency in several anatomical compartments, relatively late in its developmental trajectory in the bone marrow, i.e., during granulopoiesis.

### S1PR1^hi^ neutrophils are long-lived and exhibit enhanced mitochondrial fitness

To understand the cellular basis for S1PR1^hi^ neutrophil accumulation *in vivo* in the circulation and organs, we examined cell survival and death parameters. During sterile peritonitis, neutrophil numbers at 12 h, but not 4 h, were significantly higher in TG mice, consistent with increased neutrophil survival at later time points in this model (Fig. 3A). Splenic neutrophils exhibited a reduced frequency of Annexin V+7-AAD+ cells, indicating reduced late apoptosis/necrosis. In contrast, early apoptotic neutrophils (Annexin V^+^/7-AAD^−^) were unchanged (Fig. 3B). LDH release assays confirmed reduced cell death in bone marrow neutrophils after 24 h of *ex vivo* culture (Fig. 3C). To evaluate *in vivo* survival, we co-injected fluorescently labeled TG and control neutrophils into the peritoneum of WT mice. TG neutrophils persisted at higher levels at 24 and 48 h post-injection (Fig. 3D), indicating enhanced survival of TG neutrophils *in vivo*. Together, these data support reduced neutrophil turnover and death of S1PR1^hi^ neutrophils *in vivo*, which likely contribute to their accumulation in circulation and peripheral organs.

**Fig. 3.**
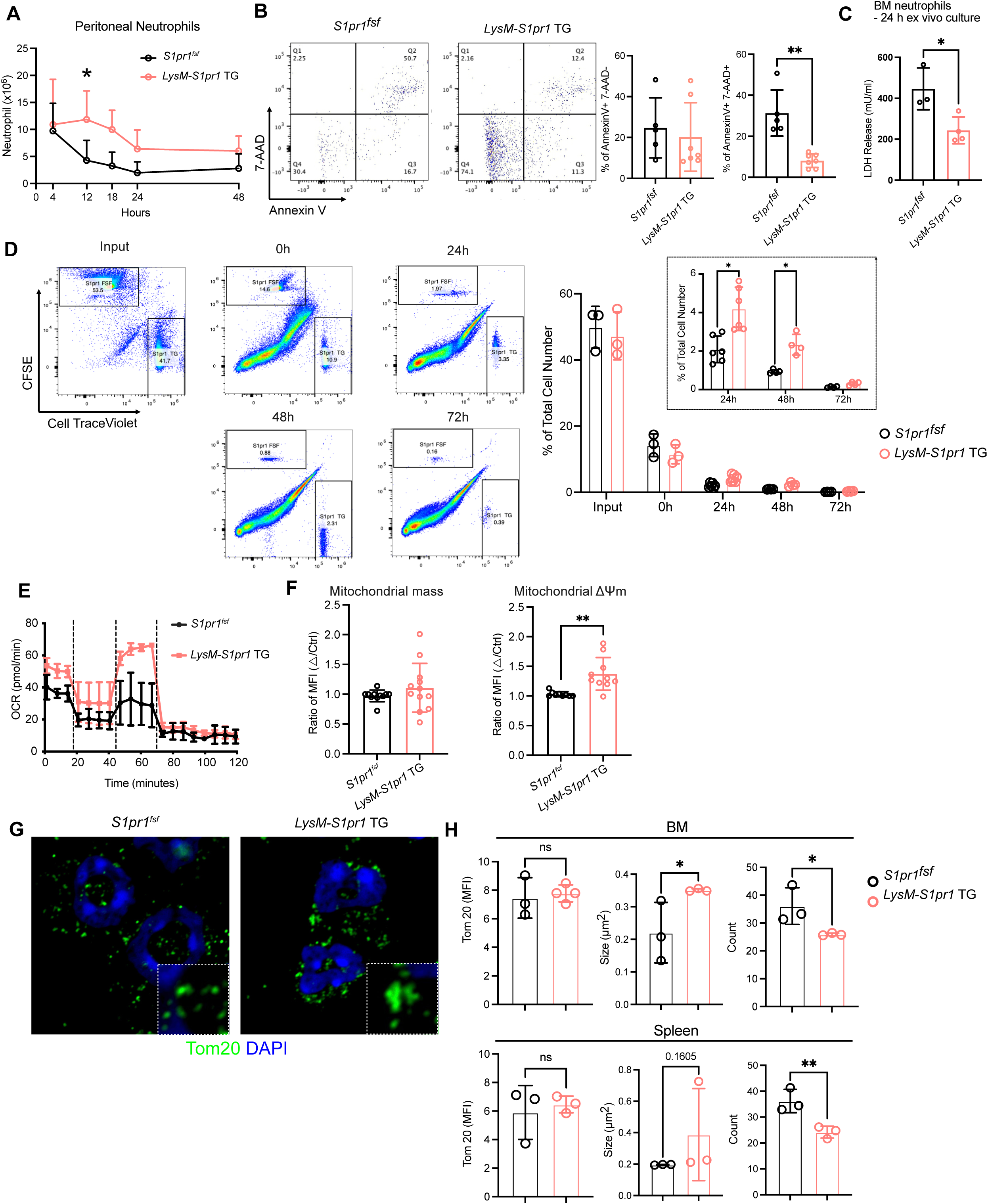
S1PR1 overexpression prolongs neutrophil survival. **(A)** Time course of peritoneal neutrophil numbers in LysM*-S1pr1* TG (n = 3) and control (n = 3) mice. **(B)** Flow cytometry of apoptotic splenic neutrophils (n ≥ 5 mice per group) stained with Annexin V and 7-AAD. Annexin V^+^7-AAD^−^cells were classified as early apoptotic, and Annexin V^+^7-AAD^+^ cells as late apoptotic/necrotic. **(C)** Lactate dehydrogenase (LDH) leakage assay was performed on bone marrow (BM) neutrophils (n ≥ 3 mice per group) after 24 h of *ex vivo* culture. **(D)** *In vivo* survival of neutrophils (n ≥ 3 mice per group) was assessed by tracking fluorescently labeled neutrophils in the peritoneal cavity at 0, 24, 48, and 72 h post-transfusion. **(E)** Oxygen consumption rate (OCR) of splenic neutrophils (n ≥ 3 mice per group) was measured by Seahorse assay. Cells (2 × 10^5^/well) were sequentially challenged with the ATP synthase inhibitor oligomycin, the uncoupler FCCP, and the complex I/II inhibitors rotenone/antimycin A. **(F)** Mitochondrial mass (left) and membrane potential (right) in splenic neutrophils (n ≥ 10 mice per group) were measured by mean fluorescence intensity (MFI) of MitoTracker Green and MitoTracker Red, respectively. **(G)** Confocal microscopy of mitochondria (Tom20, green) and nuclei (blue) in isolated splenic neutrophils. **(H)** Quantification of mitochondrial intensity, size, and count in BM and splenic neutrophils (n ≥ 3 mice per group). Data are mean ± SD. \**p* < 0.05, \*\**p* < 0.01 (two-tailed unpaired Student’s *t* test).

Since S1PR1-mediated naïve T cell survival has been linked to metabolic reprogramming (55), which involves mitochondrial content and function (40), we examined whether mitochondrial alterations in LysM*-S1pr1* TG neutrophils accompany increased persistence. Metabolic phenotyping by Seahorse analysis revealed an elevated oxygen consumption rate (OCR) in TG neutrophils under mitochondrial stress, suggesting increased oxidative phosphorylation (Fig. 3E). MitoTracker staining showed higher mitochondrial membrane potential with unchanged mass (Fig. 3F). Confocal imaging of Tom20-stained S1PR1^hi^ neutrophils revealed larger but fewer mitochondrial puncta, indicating reduced fragmentation and/or mitophagy (Fig. 3, G and H). Consistent with enhanced metabolic signaling, *S1pr1*-overexpressing BM neutrophils exhibited increased phosphorylation of mTOR downstream targets, including S6 and 4E-BP1, which was sensitive to rapamycin treatment (Fig. S4). Together, these results indicate that S1PR1 signaling enhances mitochondrial integrity and oxidative metabolism, thereby supporting metabolic fitness that promotes reduced cell death.

### S1PR1^hi^ neutrophils display transcriptional programs associated with persistence, metabolism, and reduced inflammatory signaling

Recently, single-cell transcriptomic analysis has provided a more granular understanding of neutrophil differentiation pathways that yield heterogeneous, phenotypically distinct populations (16, 17, 56, 57). To understand the transcriptional programs of S1PR1^hi^ neutrophils, we performed single-cell RNA sequencing (scRNA-seq) on peritoneal cells collected 10 hours after thioglycollate elicitation. Uniform manifold approximation and projection (UMAP) analysis identified major immune cell populations, including neutrophils, monocytes/macrophages, resident macrophages, lymphocytes (T and B), dendritic cells, mast cells, and NK cells (Fig. 4A). A representative dot plot summarizes the expression of canonical marker genes used for cluster annotation, with dot size indicating the proportion of cells expressing each gene and color intensity reflecting the average normalized expression level (Fig. 4B). Within the neutrophil compartment, UMAP projection revealed three transcriptionally distinct subsets, G5a, G5b, and G5c, as previously described (16) (Fig. 4, C and D). As reported, G5a cells abundantly expressed genes related to neutrophil migration and inflammatory responses; G5b neutrophils expressed a set of interferon-stimulated genes (ISGs); and G5c neutrophils showed high expression of genes previously associated with aging and resolution-associated states (16). We found that LysM*-S1pr1* TG mice displayed enrichment of G5c cells, consistent with enrichment of the G5c transcriptional state (Fig. 4E). These data provide a transcriptomic characterization of S1PR1^hi^ neutrophils.

**Fig. 4.**
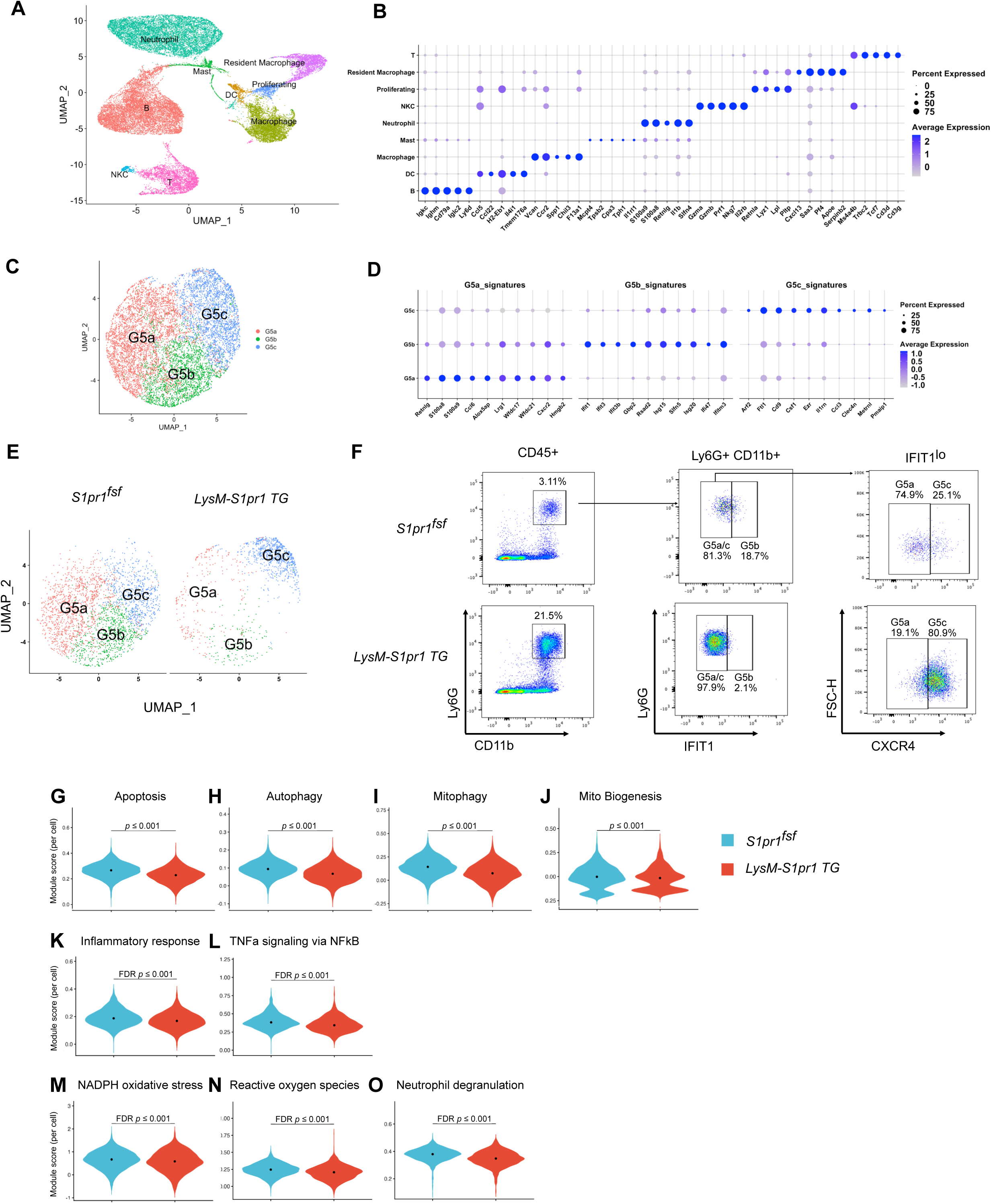
S1PR1^hi^ neutrophils exhibit a prosurvival, metabolic switch, and anti-inflammatory gene expression program. **(A)** Uniform manifold approximation and projection (UMAP) plot of peritoneal cell clusters. **(B)** Dot plot showing expression of selected marker genes across cell subsets identified by single-cell RNA sequencing. Dot size represents the percentage of cells expressing the indicated gene within each cluster, and color intensity reflects the average normalized expression level. **(C)** UMAP plot of peritoneal neutrophils clustered into G5a, G5b, and G5c. **(D)** Dot plot showing representative marker genes distinguishing G5a, G5b, and G5c neutrophil subsets. **(E)** Increased G5c and reduced G5a/b clusters in LysM*-S1pr1* TG neutrophils. **(F)** Flow cytometric validation of G5a (IFIT1^−^CXCR4^lo^), G5b (IFIT1^+^), and G5c (IFIT1^−^CXCR4^hi^) in the spleen. **(G-I)** Violin plots of per-cell module scores for apoptosis **(G)**, autophagy **(H)**, and mitophagy **(I)** in neutrophils. **(J)** Mitochondrial biogenesis module score per neutrophil. **(K-O)** Violin plots show per-cell module scores for inflammatory response **(K)**, TNFα signaling via NFkB **(L)**, NADPH oxidative stress **(M)**, reactive oxygen species **(N)**, and neutrophil degranulation **(O)**. Control cells are in blue and LysM*-S1pr1* TG cells in red. Dots mark group medians. Black brackets indicate Wilcoxon rank-sum tests performed at the cell level with Benjamini–Hochberg FDR correction; the bracket label reports the FDR p-value. Scores were computed as standardized module scores from curated gene sets; higher scores indicate stronger pathway-level expression. This cell-level analysis is exploratory.

To further validate the subcluster switch, we performed flow cytometry to distinguish splenic neutrophil populations G5a (CXCR4^lo^ IFIT1^−^), G5b (IFIT1^+^), and G5c (CXCR4^hi^ IFIT1^−^) (Fig. 4F). The results clearly documented an increase in G5c neutrophils in the spleen. These data suggested that the S1PR1^hi^ neutrophil subset resembles the G5c subset and exhibits a distinct gene expression program that correlates with increased neutrophil persistence observed *in vivo*.

The scRNAseq dataset was further analyzed using module scores with Hallmark gene sets (apoptosis) and GO/Reactome gene sets (autophagy and mitophagy). Reduced transcriptional programs for apoptosis and autophagy were observed in LysM*-S1pr1* TG neutrophils compared with control neutrophils (Fig. 4, G–I). The mitochondrial biogenesis module score per neutrophil showed distributions near zero in both groups, with a slight upward shift in LysM*-S1pr1* TG neutrophils (Fig. 4J). These results indicate that S1PR1^hi^ neutrophils exhibit transcriptional features consistent with reduced cell death signaling and suppressed mitochondrial turnover, aligning with the functional properties described above.

In addition, we noted that inflammation-and ROS-related module scores (inflammatory response, TNF_α_ signaling via NF_κ_B, NADPH oxidative stress, reactive oxygen species, neutrophil degranulation) were reduced in LysM*-S1pr1* TG neutrophils (Fig. 4, K–O), suggesting a dampened inflammatory/oxidative state. Detailed analysis of transcript expression showed reduced *Cxcr2* expression in TG neutrophils (Fig. S5A), consistent with flow cytometry data (Fig. 3E). In addition, expression of inflammation-associated genes (*Il1b*, *Casp4*, *Ptgs2*, *Nfkbia*, *Fos*, *Junb*, *Jund*, *Nfkbiz*) and cell death-associated genes (*Map1lc3b*, *Atg3*, *Stk17b*, *Ubb*, *Rb1cc1*, *Sqstm1*) was downregulated in LysM*-S1pr1* TG neutrophils (Fig. S5, B and C). Together, these findings indicate that S1PR1^hi^ neutrophils adopt a transcriptional state associated with reduced inflammatory and oxidative signaling.

### S1PR1^hi^ neutrophils retain phagocytic capacity but exhibit reduced ROS production

Phagocytosis and production of reactive oxygen species (ROS) are key effector functions of neutrophils that are essential for inflammatory, host defense, and resolution responses (58). Phagocytosis of pHrodo-*E. coli* particles was similar between LysM*-S1pr1* TG and control thioglycollate-elicited peritoneal neutrophils (Fig. 5A), indicating that enhanced S1PR1 signaling does not impair neutrophil phagocytic capacity.

**Fig. 5.**
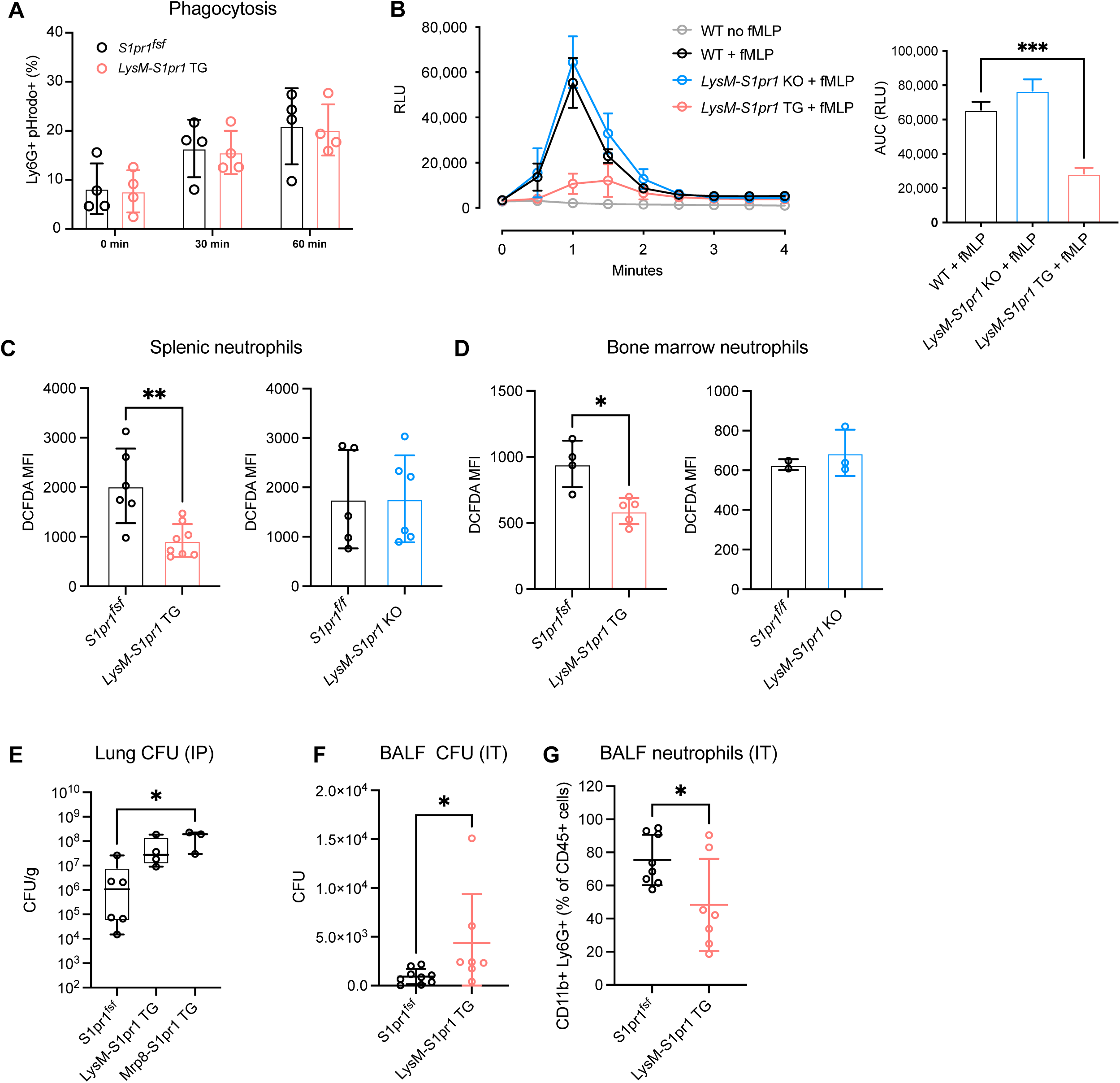
Decreased ROS production in LysM*-S1pr1* TG neutrophils. **(A)** Phagocytosis of pHrodo-*E. coli* bioparticles was assessed in thioglycollate-induced peritoneal neutrophils at 0, 30, and 60 minutes (n = 4 mice per group). pHrodo+ cells were counted as phagocytosed neutrophils, and the ratio of pHrodo+ to total neutrophils was reported. **(B)** fMLP-induced ROS were measured by luminol-based chemiluminescence (n = 3 mice per group). The area under the curve (AUC) was plotted on the right. **(C and D)** Mean fluorescence intensity (MFI) of dichlorodihydrofluorescein diacetate (DCFDA) staining for intracellular ROS in splenic **(C)** and BM neutrophils **(D)** was plotted (n ≥ 3 mice per group). (E) Bacterial burden in the lung after intraperitoneal (IP) bacterial challenge, quantified as CFU per gram of tissue. (F) Bacterial burden in bronchoalveolar lavage fluid (BALF) after intratracheal (IT) bacterial challenge, quantified as CFU. (G) Proportion of neutrophils (CD11b+Ly6G+) among CD45+ cells in BALF after IT bacterial challenge. Data are mean ± SD. \**p* < 0.05, \*\**p* < 0.01, \*\*\**p* < 0.001 (two-tailed, unpaired Student’s *t* test).

We next examined whether S1PR1 signaling affects neutrophil oxidative responses. Upon stimulation with the N-formyl peptide fMLP, activation of the NADPH oxidase complex was significantly reduced in LysM-*S1pr1* TG neutrophils, whereas responses in WT and S1PR1-deficient neutrophils were preserved (Fig. 5B). Consistent with this, intracellular ROS levels, as measured by DCFDA staining, were significantly lower in both splenic and bone marrow neutrophils from TG mice (Fig. 5, C and D). Transcriptional profiling further supported these findings, revealing reduced expression of NADPH oxidase–related genes (e.g., *Rac2*, *Ncf2*, and *Ncf4*), increased expression of antioxidant genes (e.g., *Prdx5* and *Prdx6*), and elevated Nampt, a key regulator of NAD+ biosynthesis (Fig. S6). Together, these data indicate that S1PR1 overexpression suppresses neutrophil oxidative responses. Given the central role of ROS in antimicrobial defense, we next assessed whether reduced oxidative activity affects bacterial killing. In *ex vivo* assays, extracellular bacterial killing was comparable between TG and control neutrophils. TG neutrophils exhibited a modest increase in intracellular bacterial burden at early time points, suggesting a potential delay in intracellular bacterial handling, although this difference was not statistically significant (Fig. S7). Overall, these findings indicate that reduced ROS production does not result in a major intrinsic defect in antibacterial activity under simplified *ex vivo* conditions.

To determine the functional consequences of reduced oxidative responses *in vivo*, we performed bacterial challenge experiments. After intraperitoneal (IP) bacterial challenge, LysM-*S1pr1* TG mice exhibited a higher bacterial burden in the lungs than controls (Fig. 5E). Although bacterial loads were assessed across multiple compartments, no consistent differences were observed outside the lung (Fig. S8), indicating that the defect in bacterial clearance is most prominent in this tissue. To further evaluate local antibacterial responses in the airway, mice were subjected to intratracheal (IT) bacterial challenge. TG mice displayed a significantly higher bacterial load in bronchoalveolar lavage fluid (BALF) (Fig. 5F). This was accompanied by a lower proportion of neutrophils (CD11b^+^Ly6G^+^) among CD45^+^ cells in BALF (Fig. 5G), suggesting impaired neutrophil accumulation in the airway compartment. Together, these findings indicate that reduced ROS production in S1PR1^hi^ neutrophils is associated with impaired bacterial clearance *in vivo*, with a predominant effect in the lung.

### *S1pr1* TG mice exhibit improved survival and attenuated lung injury following influenza viral infection

Neutrophil-driven inflammatory responses are crucial for viral clearance, the induction of virus-specific adaptive immunity, and the subsequent resolution and regeneration that restore homeostasis. To assess the response of myeloid S1PR1^hi^ mice to influenza infection, mice were infected intranasally with an LD_50_ dose of H1N1 virus and monitored for 21 days post-infection (dpi) (Fig. 6A). Approximately 40% of control mice died by 11 dpi, whereas only 5% of LysM*-S1pr1* TG mice died (Fig. 6B). Weight loss in LysM*-S1pr1* TG mice was significantly reduced compared with control mice (Fig. 6C). In addition, LysM*-S1pr1* TG mice exhibited better circulatory O_2_ saturation (SpO_2_) at 9 and 11 dpi (Fig. 6D), suggesting improved preservation of pulmonary function after infection. In a sublethal model (0.16 LD50) (Fig. 6E), LysM*-S1pr1* TG mice had fewer infiltrating cells in the BALF, including neutrophils and macrophages, at 3 dpi (Fig. 6, F and H). Lung viral load, as measured by nucleoprotein (NP), polymerase acidic protein (PA), and hemagglutinin (HA) gene expression, was lower in TG mice at 3 and 7 dpi (Fig. 6, I– K). BALF protein leakage and IL-6 levels were reduced at 7 dpi (Fig. 6, L and M), and IL-10 levels peaked earlier in TG mice (Fig. 6N), consistent with decreased inflammation and accelerated resolution. To determine whether these protective effects were neutrophil-intrinsic, we next evaluated influenza outcomes in Mrp8-*S1pr1* transgenic mice. Mrp8-*S1pr1* TG mice exhibited significantly improved survival, reduced weight loss, and better preservation of arterial oxygen saturation following H1N1 infection (Fig. 7, A-C). Mrp8-*S1pr1* TG mice phenocopied key protective outcomes observed in LysM-*S1pr1* TG mice, indicating that neutrophil-restricted S1PR1 overexpression is sufficient to confer protection in this model of viral pneumonia.

**Fig. 6.**
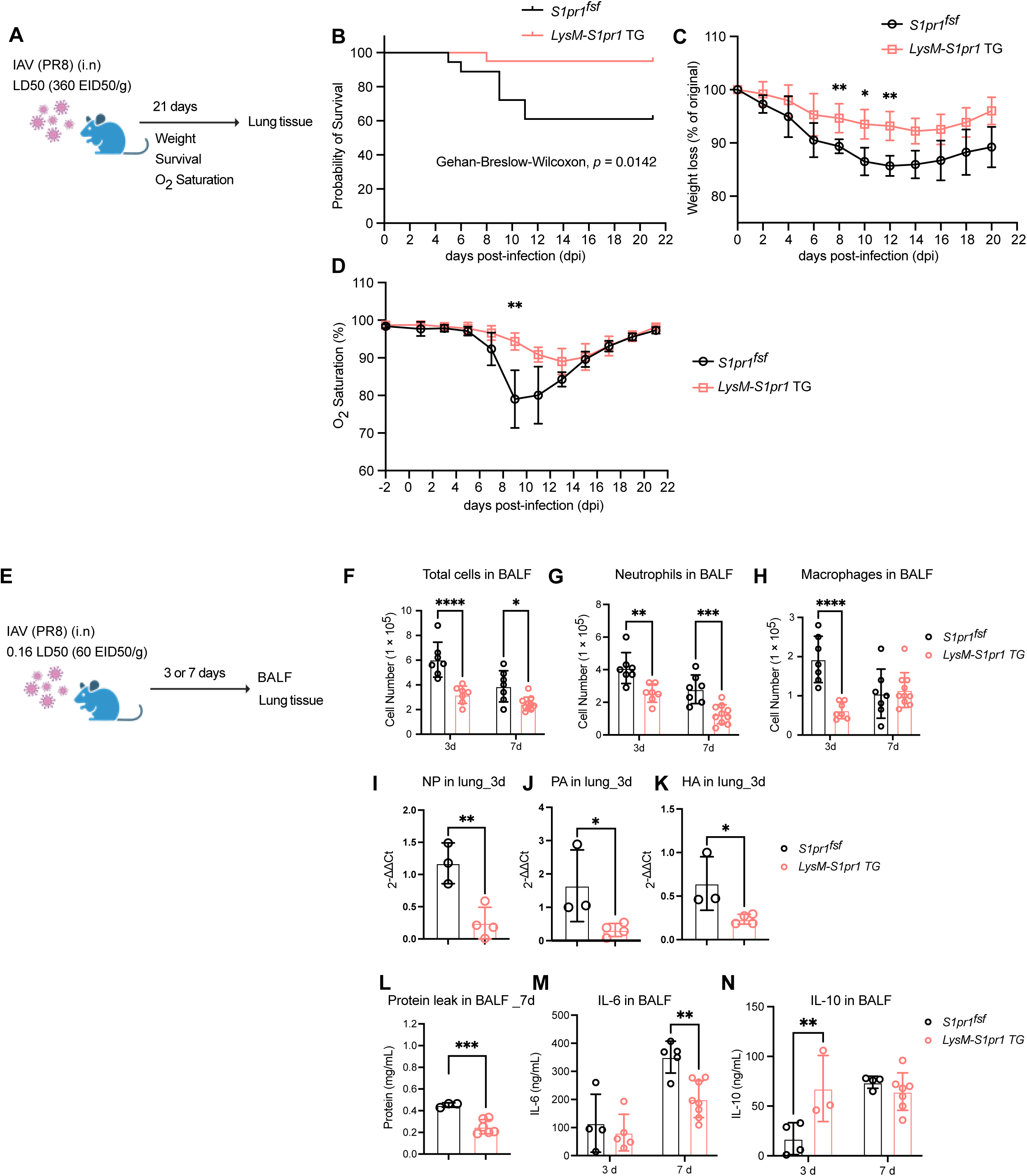
LysM*-S1pr1* TG mice are protected against H1N1-induced lung injury. **(A)** Schematic of the high-dose H1N1 (LD_50_) model over 21 days. **(B)** Survival curve of LysM*-S1pr1* TG (n = 20) and control (n = 18) mice in response to H1N1 infection. Survival curves were compared using the Gehan–Breslow–Wilcoxon test. **(C)** Body weight loss in LysM*-S1pr1* TG (n = 8) and control (n = 8) mice. **(D)** Blood oxygensaturation (SpO_2_) in LysM*-S1pr1* TG (n = 9) and control (n = 11) mice post-infection. Both (C) and (D) were analyzed by two-way mixed-effects analysis with Geisser– Greenhouse correction and Šídák’s multiple-comparison test; \**p* < 0.05, \*\**p* < 0.01. **(E)** Schematic of the low-dose (0.16 LD_50_) H1N1 model at 3 and 7 dpi. **(F)** Total cell count in BALF at 3 and 7 dpi (n = 7 mice per group). **(G and H)** BALF neutrophil and macrophage counts at 3 and 7 dpi (n = 7 mice per group). **(I-K)** Expression of viral genes, including nucleoprotein (NP), polymerase acidic protein (PA), and hemagglutinin (HA), in the infected lung in control (n = 3) and LysM*-S1pr1* TG (n = 4) mice. **(L)** Total protein in BALF in control (n = 3) and LysM*-S1pr1* TG (n = 6) mice at 7 dpi. **(M)** IL-6 (ng/mL) secretion in BALF in control (n ≥ 4) and LysM-*S1pr1* TG (n ≥ 5) mice at 3 and 7 dpi. **(N)** IL-10 (ng/mL) secretion in BALF in control (n ≥ 4) and LysM*-S1pr1* TG (n ≥ 3) mice at 3 and 7 dpi. Data are mean ± SD. \**p* < 0.05, \*\**p* < 0.01, \*\*\**p* < 0.001 (two-tailed unpaired Student’s *t* test).

**Fig. 7.**
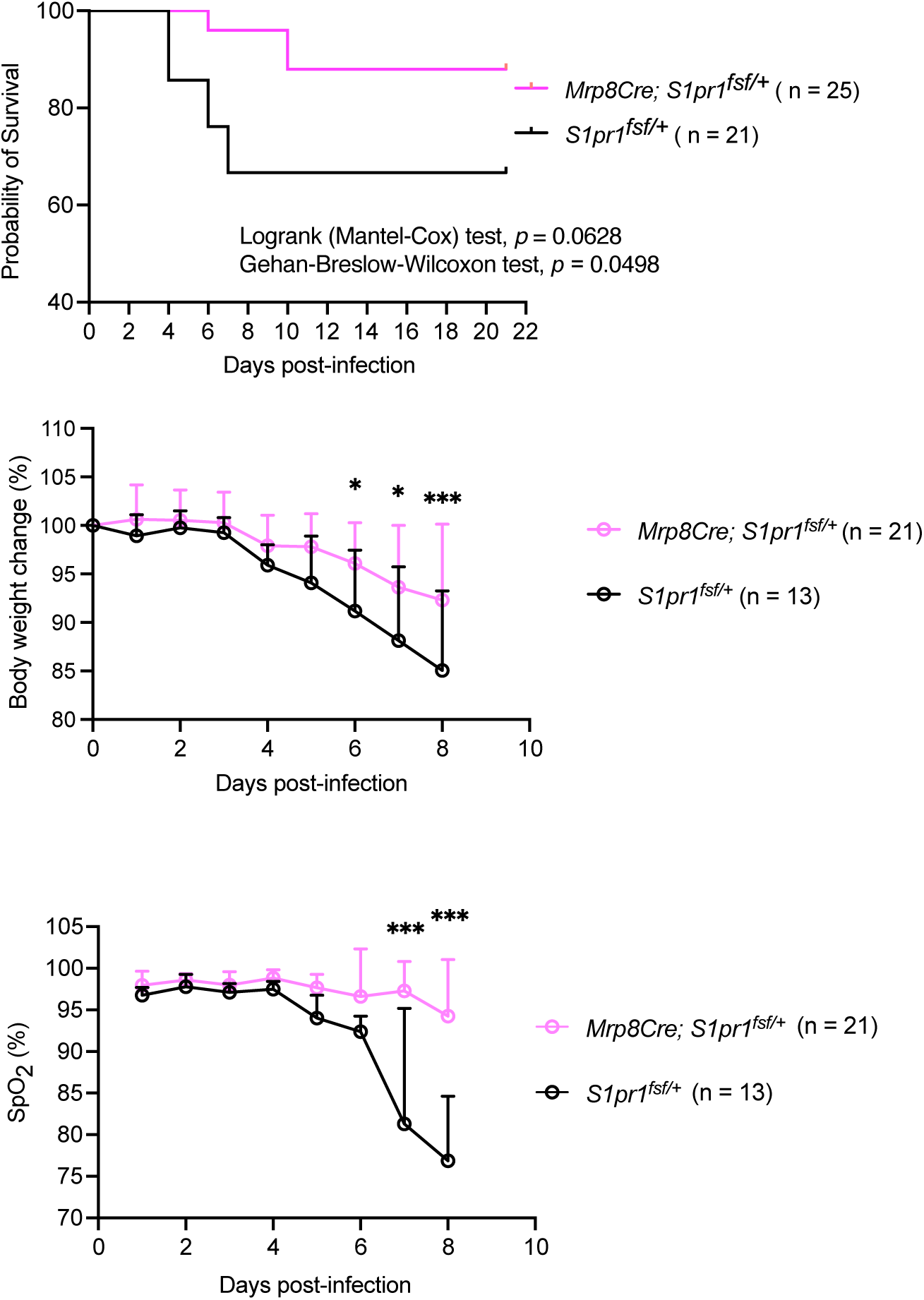
Neutrophil-intrinsic S1PR1 signaling is sufficient to improve outcomes during influenza infection. **(A)** Kaplan–Meier survival curves for Mrp8-*S1pr1* transgenic (TG) mice (n = 25) and Cre-negative littermate controls (n = 21) after intranasal infection with high-dose H1N1 (LD_50_). Survival was monitored for up to 21 days post-infection. Statistical significance was assessed using the log-rank (Mantel–Cox) and Gehan– Breslow–Wilcoxon tests. **(B)** Body weight change in control (n = 13) and Mrp8-*S1pr1* TG (n = 21) mice after influenza infection, expressed as a percentage of initial body weight. Data represent mean ± SD. (C) Arterial oxygen saturation (SpO) measured longitudinally after infection using pulse oximetry in control (n = 13) and Mrp8-*S1pr1* TG (n = 21) mice. Data represent mean ± SD. Statistical significance is denoted as \**p* < 0.05, \*\**p* < 0.01, \*\*\**p* < 0.001.

## DISCUSSION

Although S1PR1 has been extensively studied in lymphocytes and endothelial cells, its role in neutrophils remains poorly understood. Our study provides *in vivo* evidence that S1PR1 regulates neutrophil distribution, persistence, mitochondrial fitness, inflammatory phenotype, and host defense against bacteria and viruses. These effects reshape host responses in a context-dependent manner, impairing bacterial clearance while promoting protection during viral lung injury.

In this study, we used LysM*-S1pr1* KO and TG mice to define how altered S1PR1 signaling affects neutrophil biology. Unlike the TG model, LysM*-S1pr1* KO mice exhibited no discernible hematological or tissue phenotype, suggesting that S1PR1 is dispensable for neutrophil development and homeostatic functions under the steady-state conditions tested here. Although LysM-*S1pr1* KO mice showed a significant reduction in circulating lymphocyte counts, neutrophil and monocyte numbers were not significantly altered (Fig. 1A–D). In contrast, LysM*-S1pr1* TG mice displayed increased numbers of circulating and tissue-resident neutrophils, accompanied by a reduction in bone marrow neutrophils (Fig. 1). Given the established role of S1PR1 in sensing S1P gradients and regulating immune cell egress (59, 60), our findings suggest that increased S1PR1 signaling stimulates neutrophil egress from the bone marrow to peripheral tissues, including the spleen, lung, and liver. Importantly, despite increased neutrophil accumulation in peripheral tissues, S1PR1^hi^ neutrophils did not provoke overt tissue injury under homeostatic conditions.

In addition to the neutrophil phenotypes described above, LysM-*S1pr1* TG mice displayed mild macrocytic anemia, splenomegaly, and increased splenic MEP populations (Fig. S1, A-E and S3B), consistent with compensatory erythropoietic responses. FTY720 treatment normalized splenic enlargement in TG mice, supporting S1PR1–dependent splenomegaly. Notably, the anemic phenotype was not recapitulated in Mrp8-*S1pr1* TG mice (Fig. S2H). Nevertheless, Mrp8-*S1pr1* TG mice exhibited improved survival, reduced weight loss, and preserved oxygen saturation during influenza infection (Fig. 7), indicating that the protective phenotypes conferred by S1PR1 overexpression can occur independently of altered erythropoiesis.

S1PR1 is known to mediate pro-survival signaling via Gi-coupled pathways (61). Consistent with this, S1PR1-overexpressing neutrophils exhibited improved survival both *ex vivo* and *in vivo*, along with increased mitochondrial membrane potential and oxidative phosphorylation, indicative of improved mitochondrial metabolic fitness. These findings suggest that enhanced mitochondrial function supports neutrophil metabolic fitness and persistence, thereby contributing to their accumulation in LysM*-S1pr1* TG mice. Notably, S1PR1 signaling preserved mitochondrial integrity in neutrophils by reducing fragmentation and maintaining high membrane potential (Fig. 3). This observation contrasts with the well-characterized glycolytic bias of neutrophils (62–65), suggesting that S1PR1 signaling supports mitochondrial maintenance and delays mitochondrial turnover. Consistent with Gi-coupled signaling, S1PR1 may engage PI3K-mTOR pathways to support mitochondrial integrity and metabolic competence. Reduced NADPH oxidase activity and ROS output, potentially downstream of altered metabolic programming, may enhance neutrophil persistence while limiting collateral tissue damage during inflammation. Supporting this model, *S1pr1*-overexpressing neutrophils showed increased phosphorylation of mTOR downstream targets (Fig. S4), consistent with activation of metabolic signaling pathways linked to mitochondrial integrity and cell survival. Whether reduced ROS production arises directly from S1PR1-mediated signaling or secondarily from altered metabolic programming remains to be determined. While a modest alteration in *S1pr4* expression was observed in *S1pr1* transgenic neutrophils (Fig. S1G), this alteration was small and variable relative to *S1pr1* overexpression and is unlikely to be a primary driver of the observed phenotypes.

Flow cytometry and morphological analysis revealed enrichment of CXCR4^hi^CXCR2^lo^ neutrophils with band-shaped nuclei in the spleen and blood of LysM-*S1pr1* TG mice (Fig. 2), consistent with altered peripheral neutrophil states and receptor remodeling associated with trafficking. Reduced CD101 expression further supports a shift toward a non-classical neutrophil state, potentially linked to altered maturation or activation trajectories. In parallel, single-cell RNA sequencing of peritoneal neutrophils revealed an expansion of the G5c subpopulation (Fig. 4), which has been associated with neutrophil aging and reverse migration (16). In this context, our data suggest that S1PR1 signaling reprograms neutrophils toward a distinct phenotypic state characterized by enhanced persistence and reduced inflammatory output. Although we observed reduced apoptotic markers and increased neutrophil persistence in select tissues, these findings do not directly establish prolonged neutrophil lifespan but rather support a shift toward a persistence-associated neutrophil state. In addition to intrinsic state reprogramming, altered neutrophil trafficking may also contribute to these phenotypes. Neutrophils can exit tissues via lymphatic vessels and re-enter secondary lymphoid organs, a process influenced by S1P receptor signaling (66). In this framework, S1PR1 overexpression may influence the balance between tissue retention, lymphatic egress, and recirculation, thereby shaping neutrophil distribution across peripheral compartments. Consistent with this model, LysM-*S1pr1* TG mice exhibited increased neutrophil accumulation in lymphoid organs and enrichment of transcriptional features associated with aging and reverse migration. Moreover, during influenza infection, the combination of increased lung tissue neutrophils and reduced BALF neutrophils suggests altered compartmental trafficking rather than enhanced recruitment per se. Together, these findings support a model in which S1PR1 signaling reshapes neutrophil distribution and functional state through reprogramming of neutrophil phenotype, with additional contributions from altered trafficking dynamics.

Functionally, LysM*-S1pr1* TG neutrophils exhibited reduced ROS production, accompanied by downregulation of NADPH oxidase-related genes and increased expression of antioxidant programs (Fig. 5 and Fig. S6). Although phagocytosis was preserved and antibacterial activity remained largely intact under simplified *ex vivo* conditions (Fig. S7), these findings suggest there is no major intrinsic defect in bacterial killing capacity. In contrast*, in vivo* bacterial challenge revealed impaired bacterial clearance, particularly in the lung (Fig. 5E and 5F), indicating that functional defects emerge under physiological conditions. Together, these findings indicate that S1PR1 signaling dampens oxidative responses without compromising intrinsic antibacterial capacity, but instead reshapes host defense in a context-dependent manner. Reduced oxidative output may limit collateral tissue damage during inflammation, consistent with protection in viral lung injury. At the same time, impaired bacterial clearance highlights a functional trade-off associated with this reprogrammed neutrophil state, in which reduced inflammatory damage may come at the expense of optimal antibacterial defense. Future studies will be required to elucidate how S1PR1-driven neutrophil reprogramming influences host defense across diverse infectious contexts. Given the clinical use of S1PR1 modulators in autoimmune diseases, our findings suggest a potential avenue for pharmacologically tuning neutrophil function to balance antimicrobial defense and inflammatory tissue damage.

During H1N1 infection, both LysM*-S1pr1* TG and Mrp8-*S1pr1* TG mice showed improved survival, reduced weight loss, and preserved oxygen saturation (Fig. 6 and Fig. 7). Despite increased neutrophil accumulation in lung tissue, BALF neutrophils were reduced (Fig. 6), suggesting altered compartmental trafficking and reduced transmigration across the alveolar barrier rather than increased recruitment per se. Reduced viral burden, together with decreased inflammatory injury markers (BALF protein and IL-6) and altered IL-10 dynamics, supports improved resolution of lung inflammation. Together, these findings indicate that S1PR1-dependent neutrophil reprogramming promotes tissue protection by limiting inflammatory damage while preserving host control of viral infection.

Our results support a model in which elevated S1PR1 signaling reprograms neutrophils into a long-lived, low-inflammatory state marked by reduced ROS production, enhanced metabolic fitness, and altered transcriptional programs. An apparent paradox is the expansion of the G5c neutrophil cluster in *S1pr1* TG mice, previously associated with aging and apoptotic features (16). One possible explanation is that this transcriptional state reflects a late or post-maturation state rather than an irreversible commitment to cell death. In this context, S1PR1 signaling may decouple features of neutrophil aging, such as altered trafficking receptor expression and transcriptional reprogramming, from apoptotic execution, thereby promoting persistence without triggering cell death. This functional reprogramming may contribute to reduced lung damage and improved resolution during viral infection. The ability of S1PR1 to bias neutrophils toward a less-inflammatory, persistence-associated state may have therapeutic implications for inflammatory and infectious diseases. In settings of excessive neutrophil-driven inflammation, enhancing this axis may promote tissue protection and resolution. Conversely, in contexts where prolonged neutrophil persistence is detrimental, such as impaired pathogen clearance, overactivation of this pathway may be disadvantageous. Thus, the S1PR1–neutrophil axis represents a context-dependent regulatory pathway with both protective and potentially detrimental effects. Future work will be required to define how reduced ROS production influences pathogen control across different contexts. Importantly, multiple lines of evidence support a direct, cell-intrinsic role for S1PR1 in regulating neutrophil survival, metabolism, and inflammatory output.

One limitation of this study is the use of LysM-Cre, which targets multiple myeloid populations, including lymphocytes, monocytes, and macrophages. However, the core mechanistic and functional conclusions are supported by experiments in purified neutrophils, including analyses of survival, metabolic fitness, mitochondrial integrity, ROS production, and antibacterial activity, thereby reducing potential confounding effects from other cell types. Consistent with a neutrophil-intrinsic role, key phenotypes were recapitulated in neutrophil-restricted Mrp8-*S1pr1* TG mice, which exhibited improved survival, reduced weight loss, and preserved oxygen saturation during influenza infection (Fig. 7). Together, these findings support a dominant role for neutrophil-intrinsic S1PR1 signaling in the observed phenotypes.

In conclusion, our study demonstrates that neutrophil-intrinsic S1PR1 reprograms neutrophils into a persistent, metabolically fit, and low-inflammatory state. This reprogramming promotes peripheral neutrophil accumulation and reduces oxidative and inflammatory responses, thereby reshaping neutrophil function *in vivo.* Functionally, S1PR1^hi^ neutrophils exhibit context-dependent effects on host defense, with impaired bacterial clearance but enhanced protection against viral lung injury. Together, these findings support a model in which S1PR1 serves as a key regulatory axis that tunes neutrophil persistence and inflammatory output in a context-dependent manner, balancing antimicrobial defense and inflammatory tissue damage. These findings position S1PR1 as a molecular switch that uncouples neutrophil persistence from inflammatory output. These insights highlight S1PR1 as a potential target for modulating neutrophil function in infectious and inflammatory diseases.

## MATERIALS AND METHODS

### Animals

All mice were on a C57BL/6J background and used after 12 weeks of age. Both the *LysM-S1pr1* transgenic (TG) and knockout (KO) mouse lines were generated using the same myeloid-specific Cre driver line. This Cre line is commonly referred to as LysM-Cre (also known as Lyz2-Cre), in which Cre recombinase is knocked into and expressed from the endogenous lysozyme M (*Lyz2*) locus. LysM*-S1pr1* TG mice *(S1pr1^flox/stop/flox^*; LysM-Cre) were generated by crossing mice carrying a loxP-flanked transcriptional stop cassette upstream of the *S1pr1* transgene (fsf allele) with the LysM/Lyz2-Cre driver. Cre-mediated excision of the stop cassette results in enforced *S1pr1* expression in LysM-expressing myeloid cells, as previously described (67). LysM-*S1pr1* KO mice (*S1pr1^flox/flox^*; LysM-Cre) were generated by crossing *S1pr1^flox/flox^*mice with the same LysM/Lyz2-Cre driver line, resulting in myeloid-restricted deletion of the endogenous *S1pr1* allele. To assess neutrophil-intrinsic effects of S1PR1 signaling independently of broader myeloid populations, we generated Mrp8-*S1pr1* TG mice using the Mrp8-Cre (S100a8-Cre) driver line. The Mrp8-Cre transgene drives Cre recombinase expression predominantly in mature neutrophils, with minimal recombination reported in monocytes or macrophages under steady-state conditions, as previously described (68–70). Mrp8-*S1pr1* TG mice were generated by crossing Mrp8-Cre mice with *S1pr1^flox/stop/flox^* mice, resulting in enforced *S1pr1* expression preferentially in neutrophils. Unless otherwise indicated, Cre-negative littermates from the same breeding pairs were used as controls. All experimental procedures were approved by the Boston Children’s Hospital Institutional Animal Care and Use Committee.

### FTY720 treatment

Mice (n ≥ 4) were treated intraperitoneally with 0.5 mg/kg (∼150 μL per animal) of FTY720 (Sigma-Aldrich, St. Louis, MO), a functional antagonist of S1P, or vehicle (2% DMSO) every other day for 2 weeks to assess the S1PR1 dependence of splenomegaly and neutrophil distribution.

### Neutrophil isolation

Neutrophils were collected from bone marrow, spleen, or peritoneal fluid. To collect bone marrow cells, mice were euthanized with CO_2_, and their femurs and tibias were dissected and maintained in RPMI-1640 medium (Thermo Fisher, Waltham, MA). Bone marrow was exposed by clipping the ends of the bones, then centrifuged at 600 × *g* for 5 minutes, and the cell suspension was collected. To collect splenic cells, the spleen was harvested, weighed, transferred into the gentleMACS C tube (Miltenyi Biotec, Waltham, MA) containing 5 mL of RPMI-1640 medium, and dissociated in the gentleMACS dissociator (Miltenyi Biotec). RBCs were lysed with ACK lysis buffer (Thermo Fisher), and cells were washed and passed through a 40-μm cell strainer. After washing, cells were resuspended in Hanks’ balanced salt solution (HBSS) containing 0.5% fatty acid-free bovine serum albumin (ff-BSA, Sigma-Aldrich) and 2 mM EDTA. For peritoneal cells, mice were injected intraperitoneally with 2 mL of a 3% (wt/vol) thioglycollate solution (Sigma-Aldrich). Peritoneal exudate cells were collected 4 hours after thioglycollate injection by collecting lavage with 10 mL of cold HBSS. Cells were hemolyzed with ACK lysis buffer, washed, and resuspended in HBSS containing 0.5% ff-BSA and 2 mM EDTA. Cells from bone marrow, spleen, and peritoneal lavage were then purified using the Neutrophil Isolation Kit (Miltenyi Biotec) or by density gradient centrifugation with Histopaque-1119/Ficoll-Paque (71). Histopaque-1119 and Ficoll-Paque Plus were purchased from Sigma-Aldrich and GE Healthcare Life Sciences (Marlborough, MA), respectively.

### Giemsa stain

Cells in 200 μL of complete medium were loaded into an EZ Single Cytofunnel (Epredia, Kalamazoo, MI) and centrifuged at 800 rpm for 10 min to deposit onto a slide. Cells were fixed in methanol for 2 min, stained with Giemsa solution (Sigma-Aldrich) for 10 min, and rinsed with distilled water. After air-drying, cells were mounted with Neo-Mount (Sigma-Aldrich) and imaged by light microscopy. Neutrophil counts and segmented/band nuclei were quantified using ImageJ software (National Institutes of Health, Bethesda, MD).

### Gene expression analysis by qRT-PCR

Total RNA was isolated from purified neutrophils using Trizol reagent (Thermo Fisher Scientific, Waltham, MA) and the Direct-zolTM RNA MicroPrep (Zymo Research, Irvine, CA). cDNA was synthesized by reverse transcription using the qScript XLT cDNA Supermix (Quantabio, Beverly, MA). RT-PCR was performed with the PerfecTA SYBR Green Fast Mix (Quantabio) on the Step One Plus PCR system (Applied Biosystems, Foster City, CA). Relative gene expression levels were calculated using the 2-ΔΔCt method. Primers: *S1pr1*, 5’-ATGGTGTCCACTAGCATCCC-3’ (forward) and 5’-CGATGTTCAACTTGCCTGTGTAG-3’ (reverse); *Actb*, 5’-AGCCATGTACGTAGCCATCC-3’ (forward) and 5’-CTCTCAGCTGTGGTGGTGAA-3’ (reverse).

### Hematology profiling

Peripheral blood was collected from the retro-orbital cavity or the inferior vena cava and analyzed on a HEMAVET 950FS (Drew Scientific, Oxford, CT). Isolated BM and splenic neutrophils were attached to slides using a Shandon Cytospin 2 (Thermo Fisher). Blood smears or isolated neutrophils were stained with a Diff-Quick Kit (Thermo Fisher) and imaged on an Axioskop 2 mot Plus microscope (Carl Zeiss, Dublin, CA).

### Flow cytometry

Cells (1 × 10^6^) from bone marrow, spleen, liver, lung, or peritoneal lavage were surface-stained with antibodies (listed in Table S1) and analyzed by flow cytometry on a FACS Calibur or Fortessa (Becton Dickinson, Franklin Lakes, NJ) or a Cytek Aurora/Northern Light (Bethesda, MD). After gating out dead cells, neutrophils were identified as CD45^+^/CD11b^+^/Ly6G^+^ cells. Progenitor populations were defined as previously reported (60). CLPs were gated as Lin^−^IL7Rα^+^ Flt3^+^ cKit^+^ Sca-1^+^. GMPs were gated as Lin^−^ IL7Rα^−^ cKit^+^ Sca-1^−^ CD34^+^ FcγRII/III^hi^. MEPs were gated as Lin^−^ IL7Rα^−^ cKit^+^ CD34^−^ FcγRII/III^lo/-^. CMPs were gated as Lin^−^ IL7Rα^−^ cKit^+^ CD34^+^ FcγRII/III^lo/-^. For intracellular staining, surface-stained cells were fixed in 2% PFA for 10 min, washed, permeabilized in Intracellular Staining Permeabilization Wash Buffer (BioLegend, San Diego, CA), and then stained with antibodies (listed in Table S1). Fluorescence minus one (FMO) and single-stain controls were used to set gates.

### Ex vivo neutrophil survival

Splenic neutrophils were incubated at 37°C in calcium-containing HBSS for 1h in a CO_2_ incubator. After washing, cells were stained with Annexin-V and 7-AAD for 15 minutes on ice and analyzed by flow cytometry. Neutrophils were gated as CD11b^+^ Ly6G^+^ cells. Early apoptotic cells were Annexin V^+^7-AAD^−^; late apoptotic/necrotic cells were Annexin V^+^7-AAD^+^.

### In vivo neutrophil survival

Isolated BM neutrophils from LysM*-S1pr1* TG and control mice were stained with fluorescent dyes CFSE or CellTrace^TM^ Violet (1:1000 dilution) in HBSS for 20 min. After a brief wash with HBSS containing 0.5% ff-BSA, TG and control neutrophils were mixed 1:1 (∼1.4 ×10^6^ cells) and co-injected intraperitoneally into WT mice. Peritoneal cells were harvested at 0, 24, 48, and 72 hours and analyzed by flow cytometry.

### Sterile inflammation

To induce sterile inflammation, mice were injected intraperitoneally with 2 mL of 3% (wt/vol) thioglycollate solution (72). Peritoneal lavage fluids were collected 4, 12, 18, 24, or 48 hours after injection. Lavages were centrifuged to separate cells and supernatant. Cells were washed and incubated with ACK lysis buffer to remove RBCs. After washing, cells were resuspended in HBSS, and cell numbers were counted using a hemocytometer. To determine neutrophil and macrophage numbers in the cell suspension, 1 × 10^^6^ cells were incubated with Fc-blocker, followed by antibodies, and analyzed by flow cytometry. Neutrophils were gated as CD11b^+^ Ly6G^+^, and macrophages were gated as CD11b^+^ Ly6G^−^ F4/80^+^.

### ROS production

The detailed procedures (46) were followed. Briefly, peritoneal-derived neutrophils (5 × 10^5^ cells/well) were incubated in 100 μl of PBSG (PBS with glucose) containing 50 μM isoluminol and 5 U/mL horseradish peroxidase in a 96-well plate. Cell suspensions were then stimulated with 10 μM fMLP (N-formyl-Met-Leu-Phe, F3506, Sigma-Aldrich), and the plate was read on a SpectraMax M2 (Molecular Devices, San Jose, CA) over 5 minutes. The area under the curve was calculated using GraphPad Prism 7 (GraphPad Software, San Diego, CA). For DCFDA-based intracellular ROS measurement, neutrophils were stained with 10 μM DCFDA (Sigma-Aldrich) for 30 min at 37°C, washed, and analyzed by flow cytometry.

### Phagocytosis assay

Peritoneal neutrophils (2 × 10^5^ cells) were incubated in HBSS with 100 μL of the pHrodo fluorescent bioparticle conjugate (1 mg/mL, Thermo Fisher) for 30-120 minutes at 37°C. Cells with pHrodo, placed on ice, served as the control groups. After incubation for the indicated time, 2 mL of ice-cold HBSS was added to stop phagocytosis. Cells were blocked with an Fc-blocker for 10 minutes, stained with PE-Ly6G and APC-F4/80 antibodies for 30 minutes, washed, and then stained with the nucleic acid dye 7-AAD for 15 minutes. Flow cytometry analysis was performed using a FACS Calibur. The neutrophil phagocytic index was quantified using the mean fluorescence intensity (MFI) of pHrodo in 7-AAD^−^Ly6G^+^ cells.

### Neutrophil intracellular bacterial killing assay

Isolated bone marrow neutrophils were resuspended at 0.5 × 10^6^ cells/well in RPMI 1640 supplemented with 2% heat-inactivated FBS and incubated at 37 °C for 30 min before infection. *E. coli* (DH5α) was grown overnight in LB broth (Sigma-Aldrich), washed, and adjusted to an OD□□□ of 0.5 (≈ 5 × 10□ CFU/mL). Bacteria were opsonized with 10% autologous mouse serum in PBS at 37 °C for 1 h, washed, and added to neutrophil cultures at a 1:10 (neutrophil:bacterium) ratio. After incubation for 0, 30, 60, and 120 min at 37 °C, cells were centrifuged at 500 × g for 5 min, and extracellular bacteria were removed by treatment with gentamicin (250 µg/mL, Sigma-Aldrich) for 20 min, followed by three PBS washes. The final wash supernatant was plated on LB agar to confirm elimination of extracellular bacteria. Neutrophil pellets were lysed in 1 mL of 0.1% Triton X-100 for 10 min at room temperature, vortexed briefly, and then serially diluted. Diluted samples were plated on LB agar for CFU enumeration after overnight incubation at 37 °C. Bacterial suspensions incubated without neutrophils served as growth controls.

### Neutrophil extracellular bacterial killing assay

Purified neutrophils were resuspended at 0.5 × 10□ cells/well in RPMI 1640 supplemented with 2% heat-inactivated FBS and 10 µg/mL cytochalasin D (Sigma-Aldrich) to inhibit phagocytosis. Cells were incubated at 37 °C for 30 min before bacterial challenge. *E. coli* (DH5α) was cultured overnight in LB broth, washed, and adjusted to an optical density at 600 nm (OD□□□) of 0.5, corresponding to approximately 5 × 10□ CFU/mL. Neutrophils were then incubated with S. aureus at a 1:2 (neutrophil:bacterium) ratio for 0, 30, 60, and 120 min at 37 °C. At each time point, culture supernatants were serially diluted in PBS and plated onto LB agar plates, which were incubated overnight at 37 °C for colony enumeration. Bacterial suspensions incubated in medium without neutrophils served as controls to assess bacterial growth in the absence of immune cells.

### Seahorse metabolic quantification

Neutrophils (2 × 10^5^ cells/well) were resuspended in assay media and plated in microplates precoated with 22.4 μg/mL Cell-Tak (Corning) in PBS. Assay media was prepared in XF RPMI complete assay media supplemented with 10 mM glucose, 2 mM L-glutamine, and 1 mM sodium pyruvate. After incubating at 37°C without CO_2_ for 45 min, transfer the cartridge and plate to the Seahorse XFe96 Analyzer (Agilent Technologies). Oxygen consumption rate (OCR) was measured at baseline and after sequential injections of ATP synthase inhibitor oligomycin (2.5 μM, Sigma-Aldrich), uncoupler FCCP (0.61 μM, Sigma-Aldrich), and complex I/II inhibitors rotenone/antimycin A (1 μM/0.1 μM).

### Confocal microscopy

Neutrophils in 200 μL of 0.5% ff-BSA/HBSS were cytospun onto slides by centrifugation at 800 ×g for 5 min. Cells were fixed in 4% PFA for 10 min at room temperature, stained with anti-Tom20 (1:200, NBP2-67501, Novus Biologicals, Centennial, CO) overnight, and incubated with the secondary fluorescent antibody and DAPI for 1 h. Cells were then mounted with ProLong Gold. Airyscan images were acquired on a Zeiss LSM800 confocal microscope. Mitochondrial intensity, size, and count were quantified using ImageJ.

### Intraperitoneal (IP) bacterial infection

For systemic bacterial challenge, mice were infected via intraperitoneal (IP) injection with Escherichia coli (*E*. coli K12). Briefly, bacteria were cultured to mid-log phase, washed, and resuspended in sterile phosphate-buffered saline (PBS). Mice were injected intraperitoneally with 1.66 × 10^6^ colony-forming units (CFU)/g in a total volume of 200 μL. At 14 hours post-infection, mice were euthanized, and samples were collected for bacterial burden analysis. Blood was collected via cardiac puncture, and organs, including the lung, liver, and brain, along with peritoneal lavage fluid, were harvested aseptically. Tissues were homogenized in sterile PBS, serially diluted, and plated on LB agar. After overnight incubation at 37°C, CFUs were quantified. Bacterial burden was expressed as CFU per gram of tissue or per milliliter of fluid.

### Intratracheal (IT) bacterial infection

To assess local antibacterial responses in the lung, mice underwent intratracheal (IT) bacterial challenge. Mice were anesthetized with isoflurane, and E. coli K12 (3 × 10^6^ CFU in 50 μL PBS) was delivered intratracheally via a sterile catheter. At 24 hours post-infection, bronchoalveolar lavage fluid (BALF) was collected by flushing the airways with 3 mL of sterile PBS. BALF was centrifuged to separate cells and supernatant. Supernatants were used to quantify bacterial CFU and measure protein, while cell pellets were analyzed by flow cytometry. Lungs were harvested, homogenized in sterile PBS, and plated for CFU enumeration as described above. Bacterial burden was reported as CFU per lung or per milliliter of BALF. BALF cells were stained with fluorophore-conjugated antibodies against CD45, CD11b, and Ly6G to identify neutrophils (CD45^+^CD11b^+^Ly6G^+^). Data were acquired on a flow cytometer and analyzed using FlowJo software. Neutrophil frequency was expressed as a percentage of total CD45^+^ cells. Total protein concentration in BALF supernatants was measured using a bicinchoninic acid (BCA) protein assay according to the manufacturer’s instructions.

### Influenza infections and viral load quantification

The Influenza A/Puerto Rico 8/1934 (PR8) H1N1 strain was obtained from Charles River (Cat# 10100374, Wilmington, MA), aliquoted, and stored in liquid nitrogen. Mice were anesthetized i.p. with Ketamine/Xylazine (Patterson Veterinary, Devens, MA) and infected intranasally with 60 egg infectious dose (EID_50_)/gram or 0.16 lethal dose 50 (LD_50_) (73, 74) for all experiments except survival experiments. Survival experiments were performed using an LD_50_ dose, as determined by *in vivo* virus titration, at 360 EID_50_/g (73, 74). Body weight was measured every day. Viral titers were determined by quantifying viral transcripts using reverse transcription (RT)-quantitative (q) PCR, as previously described (74–77). Primers: Polymerase acidic protein gene (PA), 5’-CGGTCCAAATTCCTGCTGA-3’ (forward) and 5’-CATTGGGTTCCTTCCATCCA-3’ (reverse); Nucleoprotein (NP), 5’-CAGCCTAATCAGACCAAATG-3’ (forward) and 5’-TACCTGCTTCTCAGTTCAAG-3’ (reverse); Hemagglutinin (HA), 5’-GAGGAGCTGAGGGAGCAAT-3’ (forward) and 5’-GCCGTTACTCCGTTTGTGTT-3’ (reverse).

### Oxygen pulse meter

Oxygen saturation was measured every other day during the viral infection. Mice were shaved around the neck, where the O2 sensor was placed, and measurements were taken while the mice were awake using the MouseOx Plus (Starr Life Sciences Corp, Oakmont, PA).

### Bronchoalveolar lavage fluid (BALF) collection

The BALF collection was performed as previously described (78). Briefly, mice were euthanized by intraperitoneal injection of a lethal dose of ketamine/xylazine. The neck was disinfected with 70% ethanol, and the trachea was exposed via a midline incision. A 26-gauge needle was used to puncture the trachea between two cartilage rings, and a sterile catheter was inserted (∼0.5 cm) and secured with cotton thread. A 1 mL syringe containing 1 mL of sterile HBSS with 2 mM

EDTA was connected to the catheter. The solution was gently instilled into the lungs and aspirated while the thorax was massaged. Approximately 700–900 µL of lavage fluid was recovered per instillation, and the procedure was repeated twice. The pooled fluid was centrifuged at 400 × g for 5 min at 4 °C. Supernatants were collected for protein analysis or stored at −80 °C. Cell pellets were resuspended in 200 µL of ACK lysis buffer for 2 min at room temperature, diluted with 1 mL of ice-cold HBSS, and centrifuged again at 400 × g for 5 min. The final cell suspension was prepared in HBSS with 0.5% ff-BSA and 2 mM EDTA for flow cytometry or cytological analysis.

### Single-cell RNA sequencing (scRNA-seq) analysis

The scRNA-seq analysis was performed on peritoneal cells isolated from mice 10 h after thioglycolate injection. Approximately 1.6 × 10^9^ paired-end reads were obtained from 10× scRNA-seq using the Chromium Next GEM 3’ kit v3.1 and NovaSeq 6000 (79). Raw sequencing data were processed with CellRanger and analyzed in R (version 4.3.1) using the Seurat (version 4.0.5) package, yielding ∼33,000 cells with an average of 5,660 unique molecular identifiers (UMIs) and 1,672 genes per cell. Cell clusters were annotated using canonical markers, and neutrophil subsets (G5a/b/c) were identified as previously described (16). Differential expression was assessed using FindMarkers (log_2_ fold change > 0.25; adjusted *p* < 0.05). Module scores were computed with AddModuleScore in Seurat. For each pathway, the mean expression of its member genes (MSigDB Hallmark sets for Apoptosis; GO/Reactome sets for Autophagy and Mitophagy) was centered using matched control genes with matched baseline expression, yielding a relative activity score per cell. Violin plots display the density of cells across module scores; center lines indicate modal values without assuming a parametric distribution. Pathway-level scores were standardized module scores across curated gene sets, with higher scores indicating stronger pathway activity.

### Statistical analysis

All data, except scRNA-seq data, were analyzed in Prism (GraphPad, Boston, MA) and presented as mean ± SD. Statistical significance was assessed using one-or two-tailed unpaired Student’s t-tests or ANOVA, as appropriate. *p* values less than 0.05 were considered significant. Experiments were performed with at least 3 independent biological replicates. Survival curves in the infection model were compared using the Gehan–Breslow–Wilcoxon test (GraphPad Prism) to emphasize early differences, and similar results were obtained using the log-rank (Mantel– Cox) test. For scRNA-seq violin plots, statistical comparisons of modal values were performed using Wilcoxon rank-sum tests at the cell level, followed by Benjamini–Hochberg FDR correction. Bracket labels report FDR *p*-values. This cell-level analysis is exploratory.

## Supplementary Materials

Figs. S1 to S5

Table S1

## Funding

National Institutes of Health grants R01AI173377 and R01HL167723 to T.H., R01 HL162642 to J.O.M., and R01AI142642, R01AI145274, R01AI141386, R01HL092020, and P01HL158688 to H.R.L.

Postdoctoral Research Abroad Program, Ministry of Science and Technology, Taiwan (Y.L)

NIH training grant T32HL066987 and the Cotran-Gimbrone Research Award (A.Y.H.)

American Heart Association grant 24POST1195961 (A.G.)

AbbVie-Harvard Medical School Alliance (J.O.M.)

The Pew Charitable Trusts Biomedical Scholars (J.O.M.)

The New York Stem Cell Foundation (J.O.M.)

The Cell Discovery Network, a collaborative funded by The Manton Foundation and The Warren Alpert Foundation at Boston Children’s Hospital (J.O.M.)

Cancer Research Institute CRI Irvington postdoctoral fellowship (S.W.K.)

## Author contributions

Conceptualization: Y.L., T.S., T.H.

Methodology: Y.L., T.S., A.Y.H., A.C., A.K., M.V.L., A.G., I.F., V.A.B., S.G., R.C., J.O.M., S.W.K.

Investigation: Y.L., T.S., A.Y.H., A.C., A.K., M.V.L., A.G., I.F., V.A.B., S.G., R.C.

Visualization: Y.L., T.S.

Funding acquisition: T.H., H.R.L, J.O.M.

Project administration: T.H., Y.L., T.S.

Supervision: T.H.

Writing – original draft: Y.L., T.S., T.H.

Writing – review & editing: A.Y.H., A.C., A.K., M.V.L., A.G., I.F., V.A.B., S.G., R.C., J.O.M., S.W.K., H.R.L,

## Data and materials availability

All sequencing data generated in this study have been deposited in the Gene Expression Omnibus (GEO) under accession number GSE316812. This includes raw and processed single-cell RNA sequencing data from neutrophils isolated from LysM-*S1pr1* TG and KO mice. Additional data supporting the findings of this study are available in the article and its supplementary information files. Any further information required to reanalyze the data reported in this paper is available from the corresponding author upon reasonable request.

## References

1. Mayadas TN, Cullere X, Lowell CA. The multifaceted functions of neutrophils. Annu Rev Pathol. 2014;9:181–218.

2. Ley K, Hoffman HM, Kubes P, Cassatella MA, Zychlinsky A, Hedrick CC, et al. Neutrophils: New insights and open questions. Sci Immunol. 2018;3(30).

3. Liew PX, Kubes P. The Neutrophil’s Role During Health and Disease. Physiol Rev. 2019;99(2):1223–48.

4. Martinod K, Wagner DD. Thrombosis: tangled up in NETs. Blood. 2014;123(18):2768–76.

5. Coffelt SB, Wellenstein MD, de Visser KE. Neutrophils in cancer: neutral no more. Nat Rev Cancer. 2016;16(7):431–46.

6. McDonald B, Pittman K, Menezes GB, Hirota SA, Slaba I, Waterhouse CC, et al. Intravascular danger signals guide neutrophils to sites of sterile inflammation. Science. 2010;330(6002):362–6.

7. Rossaint J, Herter JM, Van Aken H, Napirei M, Doring Y, Weber C, et al. Synchronized integrin engagement and chemokine activation is crucial in neutrophil extracellular trap-mediated sterile inflammation. Blood. 2014;123(16):2573–84.

8. Doring Y, Soehnlein O, Weber C. Neutrophil Extracellular Traps in Atherosclerosis and Atherothrombosis. Circ Res. 2017;120(4):736–43.

9. Serhan CN, Levy BD. Resolvins in inflammation: emergence of the pro-resolving superfamily of mediators. J Clin Invest. 2018;128(7):2657–69.

10. Alnouri MW, Roquid KA, Bonnavion R, Cho H, Heering J, Kwon J, et al. SPMs exert anti-inflammatory and pro-resolving effects through positive allosteric modulation of the prostaglandin EP4 receptor. Proc Natl Acad Sci U S A. 2024;121(41):e2407130121.

11. Fredman G, Serhan CN. Specialized pro-resolving mediators in vascular inflammation and atherosclerotic cardiovascular disease. Nat Rev Cardiol. 2024;21(11):808–23.

12. Serhan CN, Chiang N. Resolvins and cysteinyl-containing pro-resolving mediators activate resolution of infectious inflammation and tissue regeneration. Prostaglandins Other Lipid Mediat. 2023;166:106718.

13. Serhan CN, Levy BD. Proresolving Lipid Mediators in the Respiratory System. Annu Rev Physiol. 2025;87(1):491–512.

14. Ghodsi A, Hidalgo A, Libreros S. Lipid mediators in neutrophil biology: inflammation, resolution and beyond. Curr Opin Hematol. 2024;31(4):175–92.

15. Ng LG, Ostuni R, Hidalgo A. Heterogeneity of neutrophils. Nat Rev Immunol. 2019;19(4):255–65.

16. Xie X, Shi Q, Wu P, Zhang X, Kambara H, Su J, et al. Single-cell transcriptome profiling reveals neutrophil heterogeneity in homeostasis and infection. Nat Immunol. 2020;21(9):1119–33.

17. Kwok I, Becht E, Xia Y, Ng M, Teh YC, Tan L, et al. Combinatorial Single-Cell Analyses of Granulocyte-Monocyte Progenitor Heterogeneity Reveals an Early Uni-potent Neutrophil Progenitor. Immunity. 2020;53(2):303–18 e5.

18. Hackert NS, Radtke FA, Exner T, Lorenz HM, Muller-Tidow C, Nigrovic PA, et al. Human and mouse neutrophils share core transcriptional programs in both homeostatic and inflamed contexts. Nat Commun. 2023;14(1):8133.

19. Garrido-Trigo A, Corraliza AM, Veny M, Dotti I, Melon-Ardanaz E, Rill A, et al. Macrophage and neutrophil heterogeneity at single-cell spatial resolution in human inflammatory bowel disease. Nat Commun. 2023;14(1):4506.

20. Palomino-Segura M, Sicilia J, Ballesteros I, Hidalgo A. Strategies of neutrophil diversification. Nat Immunol. 2023;24(4):575–84.

21. Ng LG, Ballesteros I, Cassatella MA, Egeblad M, Fridlender ZG, Gabrilovich D, et al. From complexity to consensus: A roadmap for neutrophil classification. Immunity. 2025;58(8):1890–903.

22. Soehnlein O, Lindbom L. Phagocyte partnership during the onset and resolution of inflammation. Nat Rev Immunol. 2010;10(6):427–39.

23. Calvente CJ, Tameda M, Johnson CD, Del Pilar H, Lin YC, Adronikou N, et al. Neutrophils contribute to spontaneous resolution of liver inflammation and fibrosis via microRNA-223. J Clin Invest. 2019;129(10):4091–109.

24. Horckmans M, Ring L, Duchene J, Santovito D, Schloss MJ, Drechsler M, et al. Neutrophils orchestrate post-myocardial infarction healing by polarizing macrophages towards a reparative phenotype. Eur Heart J. 2017;38(3):187–97.

25. Peiseler M, Kubes P. More friend than foe: the emerging role of neutrophils in tissue repair. J Clin Invest. 2019;129(7):2629–39.

26. Gong Y, Koh DR. Neutrophils promote inflammatory angiogenesis via release of preformed VEGF in an in vivo corneal model. Cell Tissue Res. 2010;339(2):437–48.

27. Nozawa H, Chiu C, Hanahan D. Infiltrating neutrophils mediate the initial angiogenic switch in a mouse model of multistage carcinogenesis. Proc Natl Acad Sci U S A. 2006;103(33):12493–8.

28. Cossio I, Lucas D, Hidalgo A. Neutrophils as regulators of the hematopoietic niche. Blood. 2019;133(20):2140–8.

29. Deniset JF, Surewaard BG, Lee WY, Kubes P. Splenic Ly6G(high) mature and Ly6G(int) immature neutrophils contribute to eradication of S. pneumoniae. J Exp Med. 2017;214(5):1333–50.

30. Jhunjhunwala S, Alvarez D, Aresta-DaSilva S, Tang K, Tang BC, Greiner DL, et al. Frontline Science: Splenic progenitors aid in maintaining high neutrophil numbers at sites of sterile chronic inflammation. J Leukoc Biol. 2016;100(2):253–60.

31. Koenderman L, Vrisekoop N. Extramedullary neutrophil progenitors: Quo Vadis? Cell Mol Immunol. 2024;21(8):932–4.

32. Luan Y, Hu J, Wang Q, Wang X, Li W, Qu R, et al. Wnt5 controls splenic myelopoiesis and neutrophil functional ambivalency during DSS-induced colitis. Cell Rep. 2024;43(3):113934.

33. Qu J, Jin J, Zhang M, Ng LG. Neutrophil diversity and plasticity: Implications for organ transplantation. Cell Mol Immunol. 2023;20(9):993–1001.

34. Hla T, Maciag T. An abundant transcript induced in differentiating human endothelial cells encodes a polypeptide with structural similarities to G-protein-coupled receptors. J Biol Chem. 1990;265(16):9308–13.

35. Lee MJ, Van Brocklyn JR, Thangada S, Liu CH, Hand AR, Menzeleev R, et al. Sphingosine-1-phosphate as a ligand for the G protein-coupled receptor EDG-1. Science. 1998;279(5356):1552–5.

36. Obinata H, Hla T. Sphingosine 1-phosphate and inflammation. Int Immunol. 2019;31(9):617–25.

37. Cartier A, Hla T. Sphingosine 1-phosphate: Lipid signaling in pathology and therapy. Science. 2019;366(6463).

38. Schwab SR, Cyster JG. Finding a way out: lymphocyte egress from lymphoid organs. Nat Immunol. 2007;8(12):1295–301.

39. Baeyens AAL, Schwab SR. Finding a Way Out: S1P Signaling and Immune Cell Migration. Annu Rev Immunol. 2020;38:759–84.

40. Mendoza A, Fang V, Chen C, Serasinghe M, Verma A, Muller J, et al. Lymphatic endothelial S1P promotes mitochondrial function and survival in naive T cells. Nature. 2017;546(7656):158–61.

41. Liu G, Yang K, Burns S, Shrestha S, Chi H. The S1P(1)-mTOR axis directs the reciprocal differentiation of T(H)1 and T(reg) cells. Nat Immunol. 2010;11(11):1047–56.

42. Liu G, Burns S, Huang G, Boyd K, Proia RL, Flavell RA, et al. The receptor S1P1 overrides regulatory T cell-mediated immune suppression through Akt-mTOR. Nat Immunol. 2009;10(7):769–77.

43. Garris CS, Wu L, Acharya S, Arac A, Blaho VA, Huang Y, et al. Defective sphingosine 1-phosphate receptor 1 (S1P1) phosphorylation exacerbates TH17-mediated autoimmune neuroinflammation. Nat Immunol. 2013;14(11):1166–72.

44. Finley A, Chen Z, Esposito E, Cuzzocrea S, Sabbadini R, Salvemini D. Sphingosine 1-phosphate mediates hyperalgesia via a neutrophil-dependent mechanism. PLoS One. 2013;8(1):e55255.

45. Miyabe C, Miyabe Y, Komiya T, Shioya H, Miura NN, Takahashi K, et al. A sphingosine 1-phosphate receptor agonist ameliorates animal model of vasculitis. Inflamm Res. 2017;66(4):335–40.

46. Michaud J, Kohno M, Proia RL, Hla T. Normal acute and chronic inflammatory responses in sphingosine kinase 1 knockout mice. FEBS Lett. 2006;580(19):4607–12.

47. Allende ML, Bektas M, Lee BG, Bonifacino E, Kang J, Tuymetova G, et al. Sphingosine-1-phosphate lyase deficiency produces a pro-inflammatory response while impairing neutrophil trafficking. J Biol Chem. 2011;286(9):7348–58.

48. Group CCHW. Meta-analysis of rare and common exome chip variants identifies S1PR4 and other loci influencing blood cell traits. Nat Genet. 2016;48(8):867–76.

49. Kurano M, Uranbileg B, Yatomi Y. Apolipoprotein M bound sphingosine 1-phosphate suppresses NETosis through activating S1P1 and S1P4. Biomed Pharmacother. 2023;166:115400.

50. Ye M, Iwasaki H, Laiosa CV, Stadtfeld M, Xie H, Heck S, et al. Hematopoietic stem cells expressing the myeloid lysozyme gene retain long-term, multilineage repopulation potential. Immunity. 2003;19(5):689–99.

51. Clausen BE, Burkhardt C, Reith W, Renkawitz R, Forster I. Conditional gene targeting in macrophages and granulocytes using LysMcre mice. Transgenic Res. 1999;8(4):265–77.

52. Bonecchi R, Mantovani A, Jaillon S. Chemokines as Regulators of Neutrophils: Focus on Tumors, Therapeutic Targeting, and Immunotherapy. Cancers (Basel). 2022;14(3).

53. Martelossi Cebinelli GC, de Oliveira Leandro M, Rocha Oliveira AE, Alves de Lima K, Donate PB, da Cruz Oliveira Barros C, et al. CXCR4(+) PD-L1(+) neutrophils are increased in non-survived septic mice. iScience. 2025;28(4):112083.

54. Martin C, Burdon PC, Bridger G, Gutierrez-Ramos JC, Williams TJ, Rankin SM. Chemokines acting via CXCR2 and CXCR4 control the release of neutrophils from the bone marrow and their return following senescence. Immunity. 2003;19(4):583–93.

55. Dixit D, Hallisey VM, Zhu EY, Okuniewska M, Cadwell K, Chipuk JE, et al. S1PR1 inhibition induces proapoptotic signaling in T cells and limits humoral responses within lymph nodes. J Clin Invest. 2024;134(4).

56. Evrard M, Kwok IWH, Chong SZ, Teng KWW, Becht E, Chen J, et al. Developmental Analysis of Bone Marrow Neutrophils Reveals Populations Specialized in Expansion, Trafficking, and Effector Functions. Immunity. 2018;48(2):364–79 e8.

57. Ng MSF, Kwok I, Tan L, Shi C, Cerezo-Wallis D, Tan Y, et al. Deterministic reprogramming of neutrophils within tumors. Science. 2024;383(6679):eadf6493.

58. Zhang F, Xia Y, Su J, Quan F, Zhou H, Li Q, et al. Neutrophil diversity and function in health and disease. Signal Transduct Target Ther. 2024;9(1):343.

59. Golan K, Vagima Y, Ludin A, Itkin T, Cohen-Gur S, Kalinkovich A, et al. S1P promotes murine progenitor cell egress and mobilization via S1P1-mediated ROS signaling and SDF-1 release. Blood. 2012;119(11):2478–88.

60. Blaho VA, Galvani S, Engelbrecht E, Liu C, Swendeman SL, Kono M, et al. HDL-bound sphingosine-1-phosphate restrains lymphopoiesis and neuroinflammation. Nature. 2015;523(7560):342–6.

61. Rutherford C, Childs S, Ohotski J, McGlynn L, Riddick M, MacFarlane S, et al. Regulation of cell survival by sphingosine-1-phosphate receptor S1P1 via reciprocal ERK-dependent suppression of Bim and PI-3-kinase/protein kinase C-mediated upregulation of Mcl-1. Cell Death Dis. 2013;4(11):e927.

62. Injarabian L, Devin A, Ransac S, Marteyn BS. Neutrophil Metabolic Shift during their Lifecycle: Impact on their Survival and Activation. Int J Mol Sci. 2019;21(1).

63. Jeon JH, Hong CW, Kim EY, Lee JM. Current Understanding on the Metabolism of Neutrophils. Immune Netw. 2020;20(6):e46.

64. Toller-Kawahisa JE, Hiroki CH, Silva CMS, Nascimento DC, Publio GA, Martins TV, et al. The metabolic function of pyruvate kinase M2 regulates reactive oxygen species production and microbial killing by neutrophils. Nat Commun. 2023;14(1):4280.

65. Yipeng Z, Chao C, Ranran L, Tingting P, Hongping Q. Metabolism: a potential regulator of neutrophil fate. Front Immunol. 2024;15:1500676.

66. Bogoslowski A, Wijeyesinghe S, Lee WY, Chen CS, Alanani S, Jenne C, et al. Neutrophils Recirculate through Lymph Nodes to Survey Tissues for Pathogens. J Immunol. 2020;204(9):2552–61.

67. Jung B, Obinata H, Galvani S, Mendelson K, Ding BS, Skoura A, et al. Flow-regulated endothelial S1P receptor-1 signaling sustains vascular development. Dev Cell. 2012;23(3):600–10.

68. Stackowicz J, Gaudenzio N, Serhan N, Conde E, Godon O, Marichal T, et al. Neutrophil-specific gain-of-function mutations in Nlrp3 promote development of cryopyrin-associated periodic syndrome. J Exp Med. 2021;218(10).

69. Reber LL, Gillis CM, Starkl P, Jonsson F, Sibilano R, Marichal T, et al. Neutrophil myeloperoxidase diminishes the toxic effects and mortality induced by lipopolysaccharide. J Exp Med. 2017;214(5):1249–58.

70. Wang G, Zhang C, Kambara H, Dambrot C, Xie X, Zhao L, et al. Identification of the Transgene Integration Site and Host Genome Changes in MRP8-Cre/ires-EGFP Transgenic Mice by Targeted Locus Amplification. Front Immunol. 2022;13:875991.

71. Wishart AL, Swamydas M, Lionakis MS. Isolation of Mouse Neutrophils. Curr Protoc. 2023;3(9):e879.

72. Schneider M. Collecting resident or thioglycollate-elicited peritoneal macrophages. Methods Mol Biol. 2013;1031:37–40.

73. Galani IE, Triantafyllia V, Eleminiadou EE, Andreakos E. Protocol for influenza A virus infection of mice and viral load determination. STAR Protoc. 2022;3(1):101151.

74. Griffith JW, Faustino LD, Cottrell VI, Nepal K, Hariri LP, Chiu RS, et al. Regulatory T cell-derived IL-1Ra suppresses the innate response to respiratory viral infection. Nat Immunol. 2023;24(12):2091–107.

75. Moltedo B, Li W, Yount JS, Moran TM. Unique type I interferon responses determine the functional fate of migratory lung dendritic cells during influenza virus infection. PLoS pathogens. 2011;7(11):e1002345.

76. Cho JL, Roche MI, Sandall B, Brass AL, Seed B, Xavier RJ, et al. Enhanced Tim3 activity improves survival after influenza infection. Journal of immunology. 2012;189(6):2879–89.

77. Glenn JD, Smith MD, Xue P, Chan-Li Y, Collins S, Calabresi PA, et al. CNS-targeted autoimmunity leads to increased influenza mortality in mice. J Exp Med. 2017;214(2):297–307.

78. Van Hoecke L, Job ER, Saelens X, Roose K. Bronchoalveolar Lavage of Murine Lungs to Analyze Inflammatory Cell Infiltration. J Vis Exp. 2017(123).

79. Engelbrecht E, Levesque MV, He L, Vanlandewijck M, Nitzsche A, Niazi H, et al. Sphingosine 1-phosphate-regulated transcriptomes in heterogenous arterial and lymphatic endothelium of the aorta. Elife. 2020;9.

