## Supplemental Figures and Table for "Sphingosine 1-phosphate receptor-1 signaling enhances neutrophil survival while suppressing inflammation and bacterial host defense functions": S. Figures.pdf

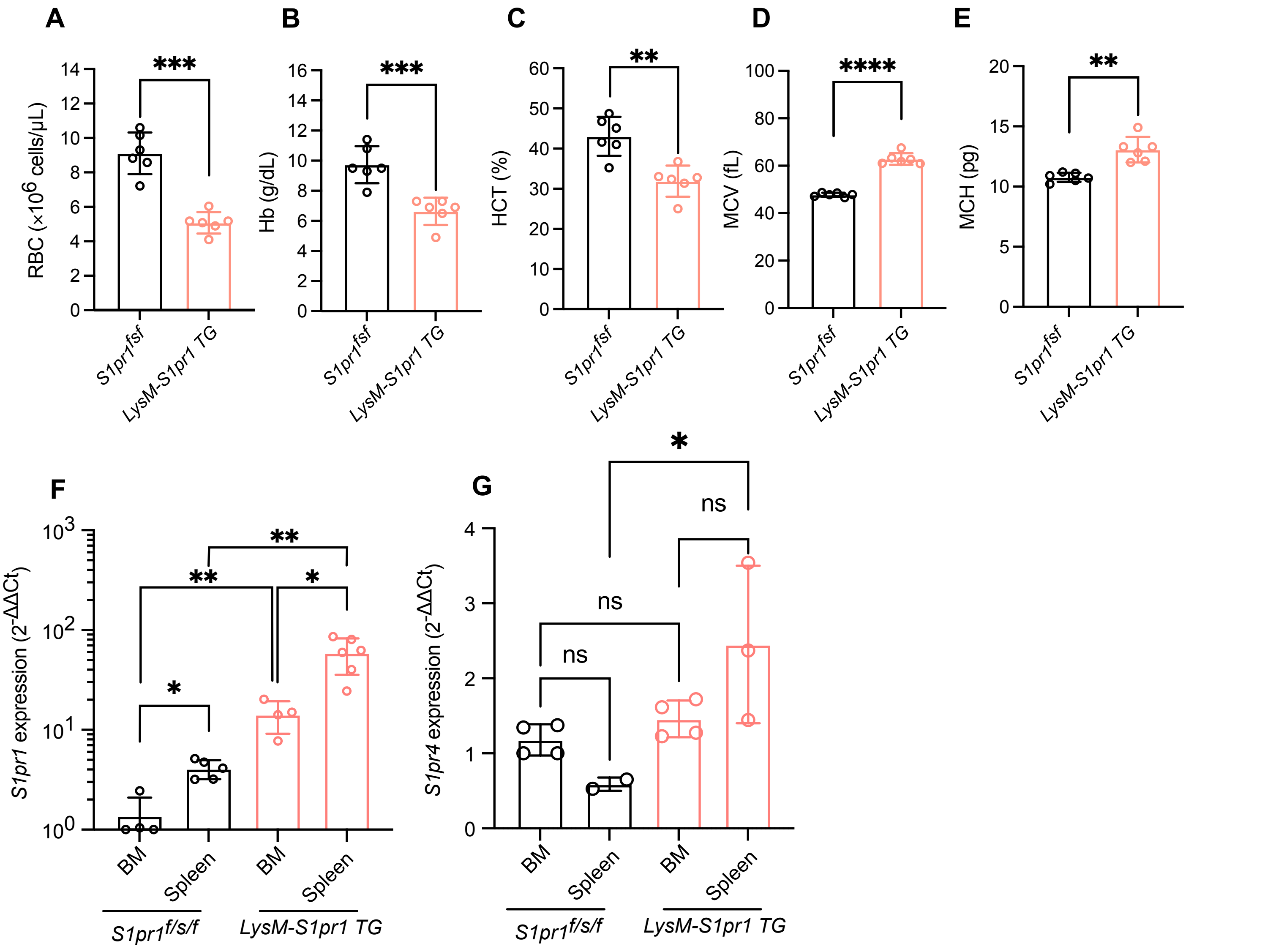


**Supplementary Figure 1. Hematologic alterations and tissue-specific S1pr1 expression in LysM-*S1pr1* transgenic mice.**
**(A-E)** Counts of red blood cells (RBCs), hemoglobin (Hb), and hematocrit (HCT), mean corpuscular volume (MCV), and mean corpuscular hemoglobin (MCH) in peripheral blood from *LysM-S1pr1* TG (n = 6) and $S1pr1^{fsf}$ (n = 6) mice. **(F and G)** qRT-PCR showed *S1pr1* and *S1pr4* expression (normalized to β-actin) in isolated BM and spleen neutrophils. Data are mean ± S.D. *$p < 0.05$, **$p < 0.01$, ***$p < 0.001$, ****$p < 0.0001$ (two-tailed unpaired Student's *t* test for blood count and Mann–Whitney U test for *S1pr1* expression).

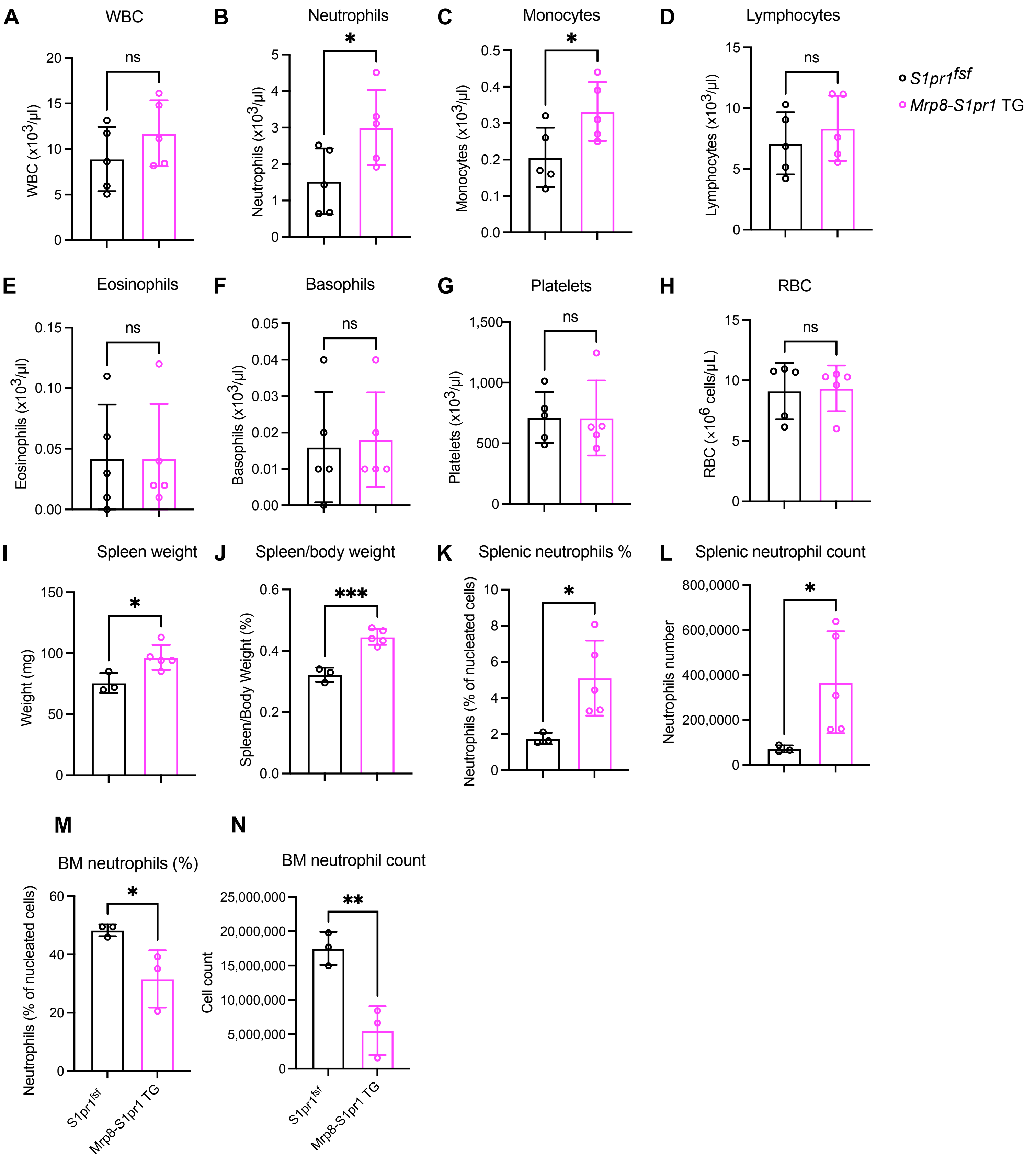


**Supplementary Figure 2. Neutrophil-specific S1pr1 overexpression increases splenic neutrophils and reduces bone marrow neutrophil retention**

**(A-H)** Counts of WBCs, neutrophils, monocytes, lymphocytes, eosinophils, basophils, platelets, and RBCs in peripheral blood from *Mrp8-S1pr1* TG (n = 5) and *S1pr1*$^{fsf}$(n = 5) mice. **(I and J)** Spleen weights and Spleen/body weight ratio of *S1pr1*$^{fsf/+}$ (n = 3) and *Mrp8-S1pr1* TG (n = 5) mice. **(K and L)** Splenic neutrophils % (K) and count (L) in *Mrp8-S1pr1* TG (n = 3) and *S1pr1*$^{fsf}$ (n = 5) mice. **(M and N)** Bone marrow (BM) neutrophil % (M) and counts (N) in *Mrp8-S1pr1* TG and *S1pr1*$^{fsf}$ mice. Data are mean ± S.D. * $p$<0.05, **$p$<0.01, *** $p$<0.001 (two-tailed unpaired Student's $t$ test).

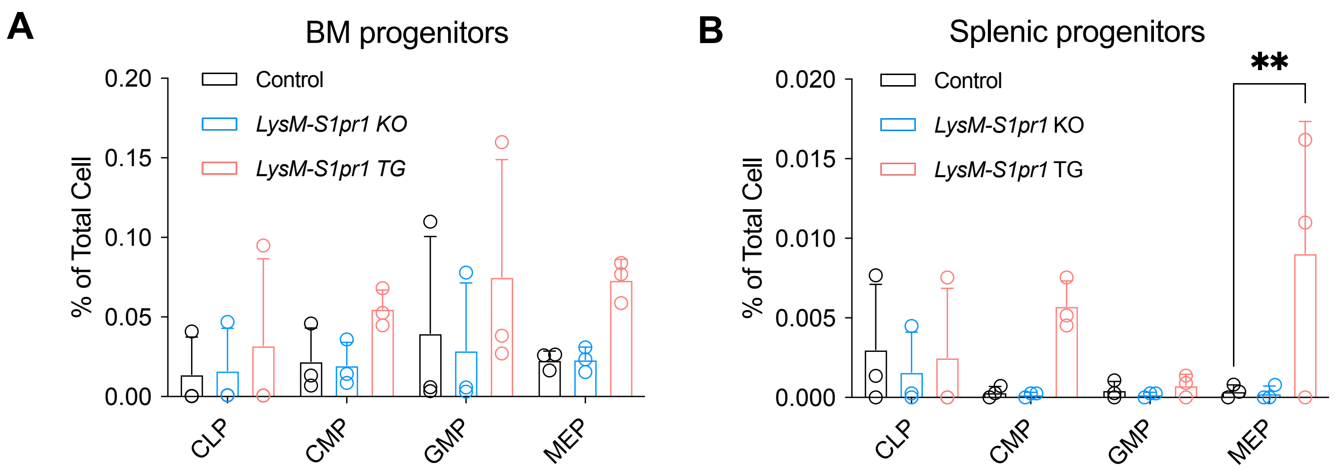


**Supplementary Figure 3. Extramedullary enrichment of megakaryocyte/erythroid progenitors in LysM-S1pr1 transgenic mice**
**(A-B)** Myeloid progenitor profile in spleen and bone marrow by flow cytometry. Common lymphoid progenitor cells, common myeloid progenitors (CMPs), granulocyte myeloid progenitors (GMPs) and megakaryocyte/erythrocyte progenitors (MEPs) were shown. Data are mean ± S.D. **$p$<0.01 (two-tailed unpaired Student's $t$ test).

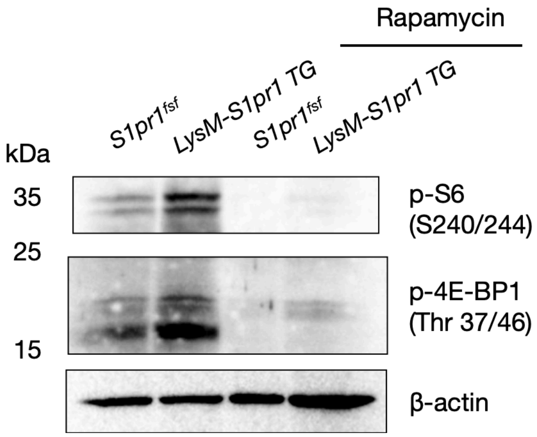


**Supplementary Figure 4. Enhanced mTOR pathway activation in *S1pr1*-overexpressing BM neutrophils.** Representative immunoblot analysis of mTOR pathway components in bone marrow (BM) neutrophils isolated from control and LysM-*S1pr1* transgenic mice. Isolated neutrophils were treated with rapamycin (50 nM) or vehicle (EtOH) for 60 min prior to lysis. Phosphorylation of downstream mTOR targets (p-S6 and p-4E-BP1) was increased in *S1pr1*-overexpressing neutrophils compared with controls, while total protein levels and loading controls (β-actin) remained comparable across samples.

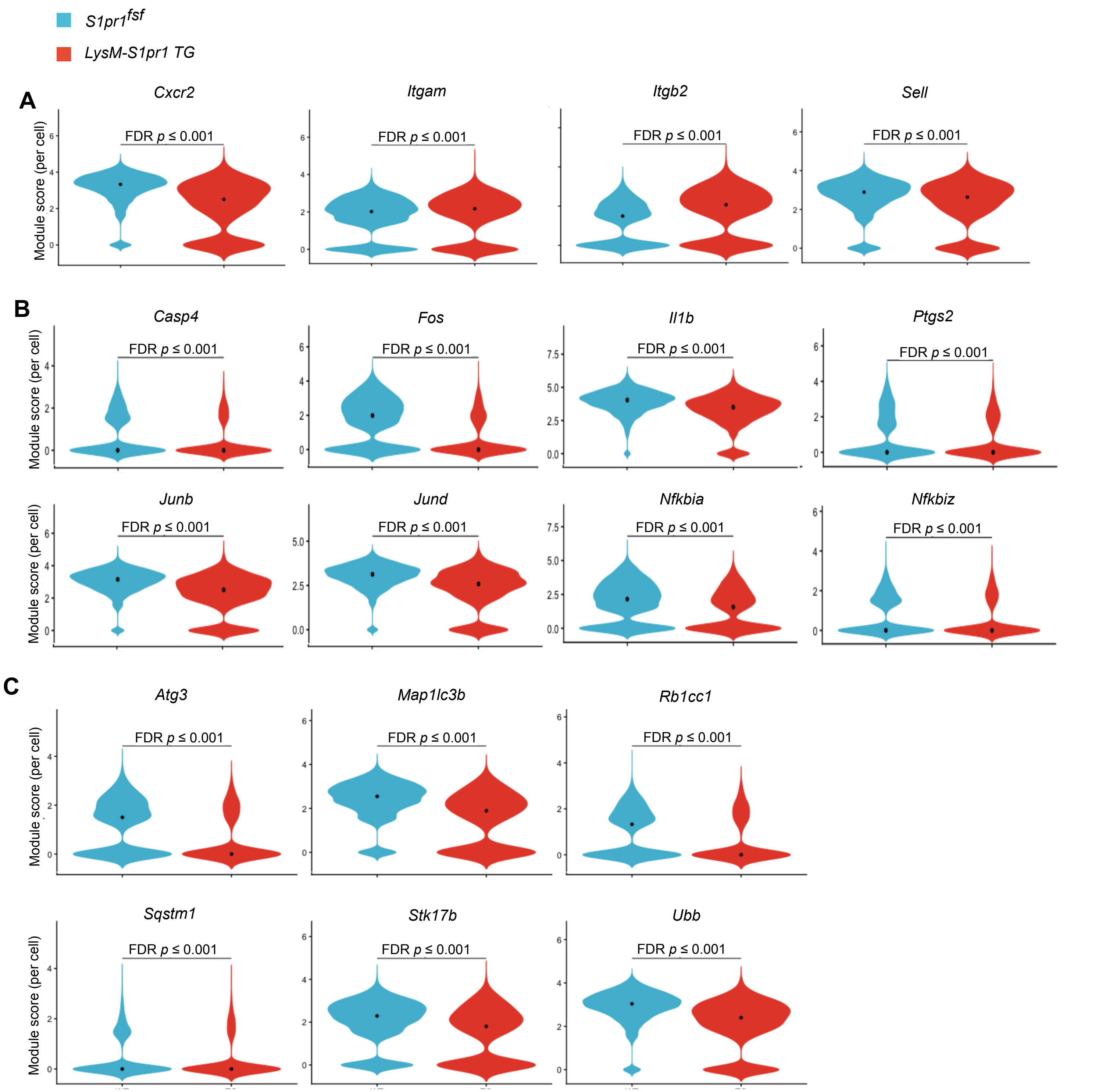


**Supplementary Figure 5. S1PR1 overexpression reshapes neutrophil transcriptional programs associated with trafficking, inflammation, and cell death.**
**(A-C)** Violin plots of trafficking **(A)**, inflammation **(B)**, and cell death genes **(C)** between *LysM-S1pr1* TG and control neutrophils. Values are scaled per gene across neutrophil clusters Wilcoxon rank-sum tests was performed at the cell level, followed by Benjamini–Hochberg FDR correction.
Bracket label reports FDR *p*-value.

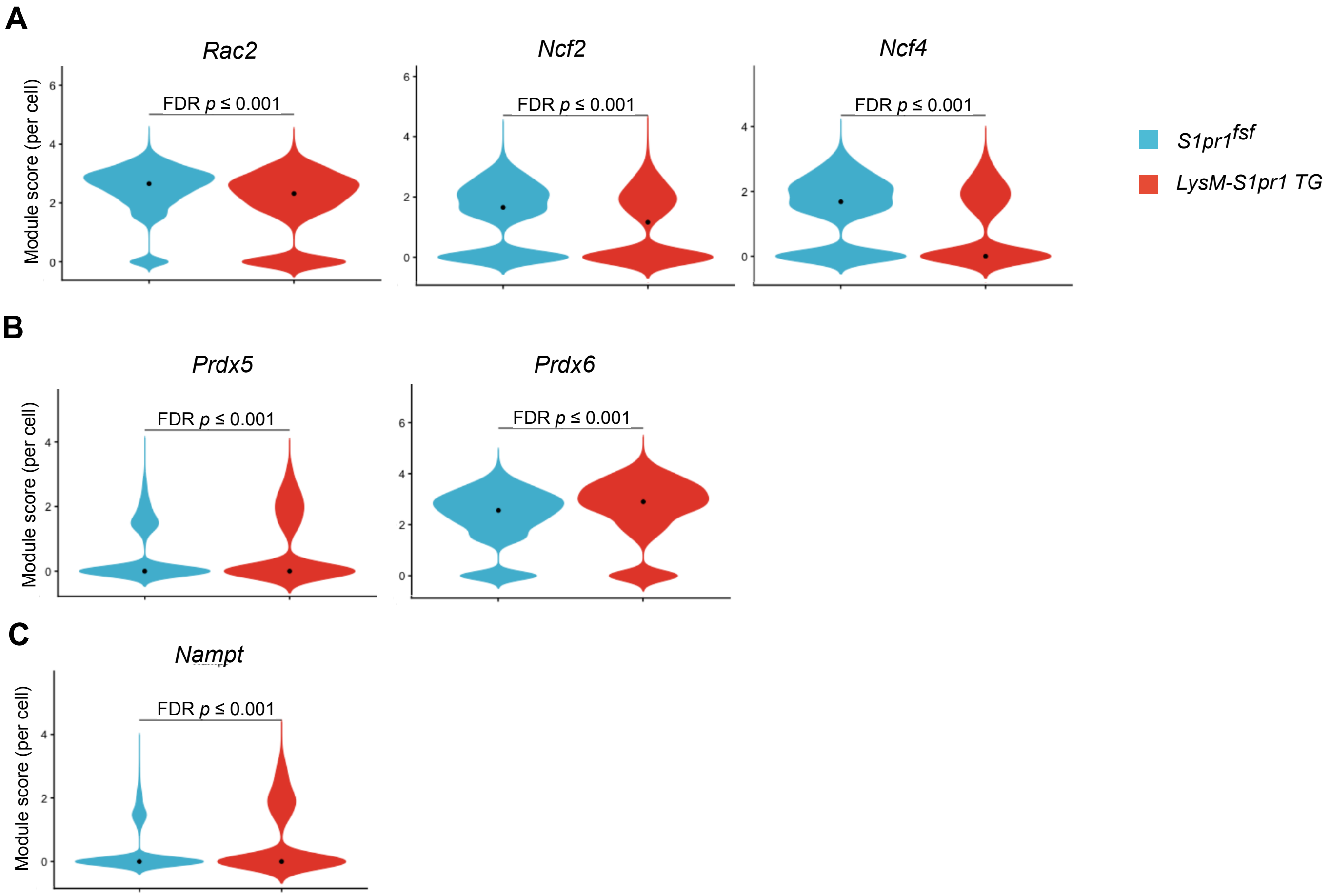


**Supplementary Figure 6. Transcriptional changes in S1PR1$^{hi}$ neutrophils related to oxidative and metabolic pathways.**
**(A)** Violin plots showing expression of NADPH oxidase–related genes (*Rac2*, *Ncf2*, and *Ncf4*) in neutrophils from *LysM-S1pr1* TG and control mice. **(B)** Expression of antioxidant genes (*Prdx5* and *Prdx6*) in neutrophils from *LysM-S1pr1* TG and control mice. **(C)** Expression of *Nampt*, a key regulator of NAD+ biosynthesis, in neutrophils from *LysM-S1pr1* TG and control mice. Expression levels are shown as module scores per cell. Statistical significance was determined using differential expression analysis with false discovery rate (FDR) correction (FDR $p \leq 0.001$).

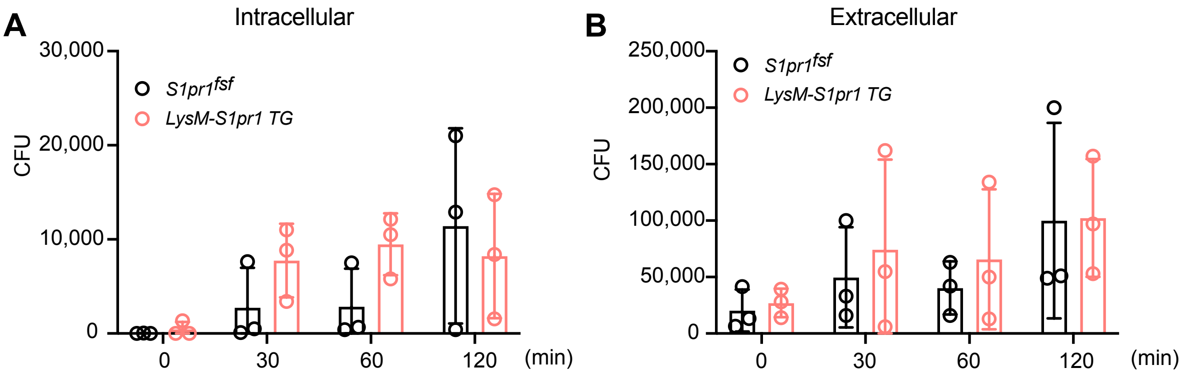


**Supplementary Figure 7. Intracellular and extracellular bacterial killing by S1PR1-overexpressing neutrophils.**

**(A-B)** Intracellular **(A)** and extracellular **(B)** *E. coli* killing assay using bone marrow neutrophils from *LysM-S1pr1* TG and control mice. For intracellular killing, neutrophils ($5 \times 10^5$) were incubated with opsonized *E. coli* K12 ($5 \times 10^6$) for 0, 30, 60, or 120 min at 37°C. Cells were treated with gentamicin (250 μg/mL) for 30 min to remove extracellular bacteria. After washing, cells were lysed with 0.1% Triton X-100, and then plated on LB agar for colony-forming unit (CFU) counts. For extracellular killing, neutrophils ($5 \times 10^5$) were incubated with *E. coli* ($1 \times 10^6$) and actin polymerization inhibitor cytochalasin D (10 μg/mL) for 0, 30, 60, or 120 min at 37°C. After centrifugation, supernatants were collected and plated on LB agar for CFU counts. A modest increase in intracellular CFU at early time points was observed in TG neutrophils, whereas extracellular killing was comparable between genotypes.

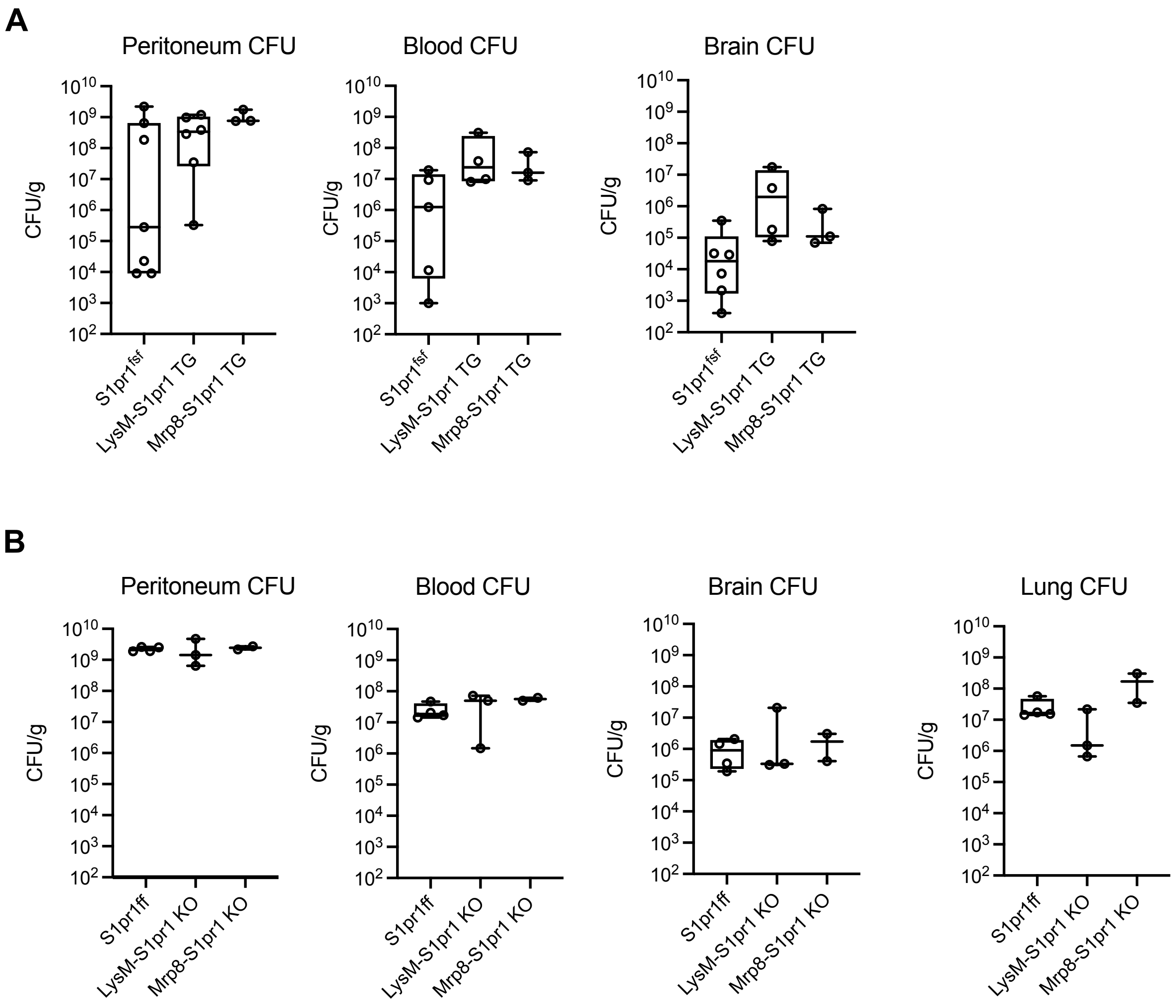


**Supplementary Figure 8. Bacterial burden across multiple compartments following intraperitoneal challenge.** Bacterial load was quantified as colony-forming units (CFU) in peritoneal fluid, blood, brain, and lung following intraperitoneal (IP) bacterial challenge. **(A)** CFU measurements in *S1pr1*$^{fsf}$ control, *LysM-S1pr1* TG, and *Mrp8-S1pr1* TG mice. **(B)** CFU measurements in *S1pr1*$^{fl/fl}$ control, *LysM-S1pr1* KO, and *Mrp8-S1pr1* KO mice. Each dot represents an individual mouse. Data are presented as mean ± SD.
