## Supplemental Figures and Table for "Sphingosine 1-phosphate receptor-1 signaling enhances neutrophil survival while suppressing inflammation and bacterial host defense functions": Table S1.pdf

**Table S1 Antibody List**

| Antibody | Fluorophore | Company | Catalog # | Clone | Dilution |
|---|---|---|---|---|---|
| AnnexinV | AF647 | BioLegend | 640912 | | |
| c-Kit (CD117) | PerCP | BioLegend | 105821 | 2B8 | 1:100 |
| CD101 | PE/C7 | ThermoFisher | 25-1011-82 | Moushi101 | 1:100 |
| CD11b | BV510 | BioLegend | 101263 | M1/70 | 1:100 |
| CD11b | FITC | BioLegend | 101205 | M1/70 | 1:100 |
| CD127 (IL7Rα) | FITC | Invitrogen | 11-1271-81 | A7R34 | 1:100 |
| CD135 (FLT3) | PE | BioLegend | 135305 | A2F10 | 1:100 |
| CD16/32 (FcγRII/III) | FITC | BioLegend | 101305 | 93 | 1:100 |
| CD16/32 (FcγRII/III) | AF700 | Invitrogen | 56-0161-80 | 93 | 1:100 |
| CD16/32 (FcγRII/III) | N/A | BD Pharmigen | 553141 | 8D 2.4G2 | 1:100 |
| CD34 | AF647 | BD Biosciences | 560233 | RAM34 | 1:100 |
| CD34 | APC/Cy7 | BioLegend | 128621 | HM34 | 1:100 |
| CD45 | BV510 | BioLegend | 103138 | 30-F11 | 1:100 |
| CD45 | APC | BioLegend | 103112 | 30-F11 | 1:100 |
| CD62L | APC | BioLegend | 104412 | MEL-14 | 1:100 |
| CXCR2 (CD182) | APC | BioLegend | 149312 | SA044G4 | 1:100 |
| CXCR2 (CD182) | APC/Cy7 | BioLegend | 149314 | SA044G4 | 1:100 |
| CXCR4 (CD184) | PerCP/Cy5.5 | BioLegend | 146510 | L276F12 | 1:100 |
| CXCR4 (CD184) | APC | BioLegend | 146507 | L276F12 | 1:100 |
| F4/80 | APC | BioLegend | 123115 | BM8 | 1:100 |
| F4/80 | APC/Cy7 | BioLegend | 123118 | BM8 | 1:100 |
| Lin | eF450 | Invitrogen | 888-7772-72 | | 1:100 |
| Ly6G | PE | BioLegend | 127608 | 1A8 | 1:100 |
| Ly6G | APC | BioLegend | 127614 | 1A8 | 1:100 |
| IFIT1 (p56) | N/A | Sigma-Aldrich | ABF117 | | 1:100 |
| S1P1 | APC | R&D system | FAB7089A | 713412 | 1:100 |
| Sca-1 (Ly-6A/E) | PE/Cy7 | BioLegend | 108113 | D7 | 1:100 |
